# From Pigs to Silkworms: Cognition and Welfare across 10 Farmed Taxa

**DOI:** 10.1101/2022.11.11.516141

**Authors:** Rachael Miller, Martina Schiestl, Anna Trevarthen, Leigh Gaffney, J. Michelle Lavery, Bob Fischer, Alexandra Schnell

## Abstract

Billions of animals across many taxa are extensively farmed, with critical impacts on animal welfare. Societal efforts to reduce animal suffering lack rigorous and systematic approaches that facilitate maximising welfare improvements, such as informed funding allocation decisions. We present a multi-measure, cross-taxa framework for modelling differences in pain, suffering, and related cognition to assess whether certain animals have larger welfare ranges (how well or badly animals can fare). Measures include behavioural flexibility, cognitive sophistication, and general learning. We evaluated 90 empirically detectable proxies for cognition and welfare range (henceforth ‘proxies’) in pigs, chickens, carp, salmon, octopus, shrimp, crabs, crayfish, bees, and silkworms. We grouped a subset of proxies into: A) 10 ideal proxies and B) 10 less ideal proxies but with sufficient data for interspecies comparisons. We graded the strength of evidence per proxy across taxa, and constructed a cognition and welfare range profile, with overall judgement scores (ranging from likely no/low confidence to yes/very high confidence). We discuss the implications of comparisons and highlight key avenues for future research. This work is timely, given recent indications of significant political will towards reducing animal suffering, such as the inclusion of cephalopods and decapods in the Animal Welfare (Sentience) Bill following a UK government-commissioned research review. Given the novelty and robustness of our review, we believe it sets a new standard for investigating interspecies comparisons of cognition and welfare ranges and helps inform future research. This should help streamline funding allocations and improve the welfare of millions of farmed animals.

**Graphical/ Visual Abstract and Caption:** Cognition and welfare in farmed animals - from pigs to silkworms (Free stock images: http://www.pixabay.com)

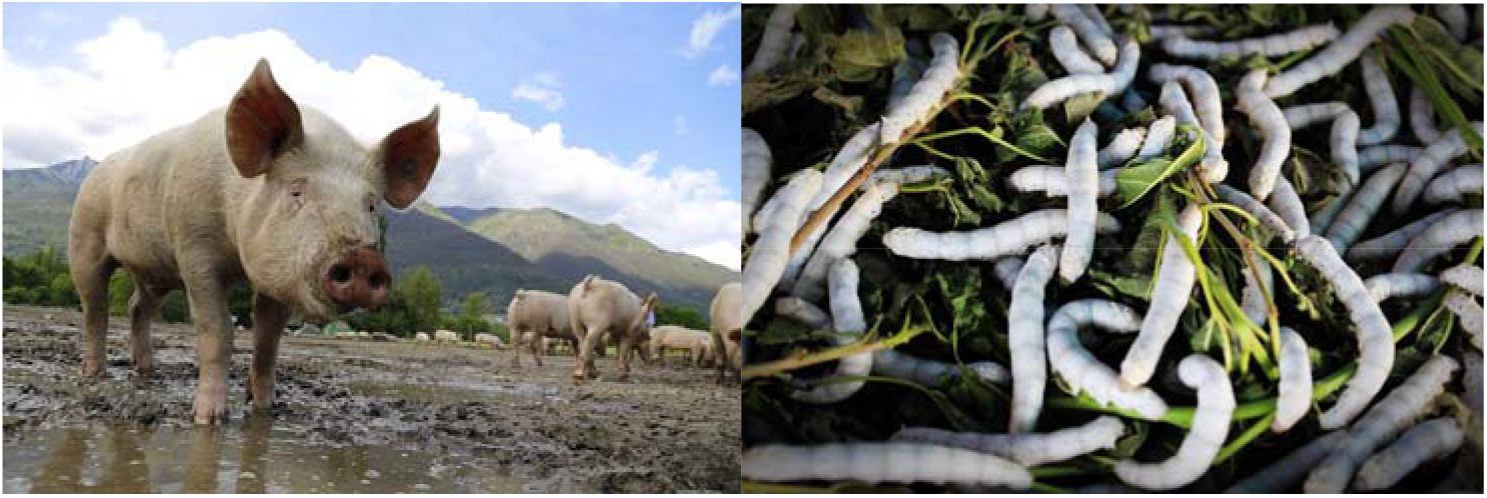

## 1. INTRODUCTION

Do certain animals have a greater capacity for suffering? This article presents and applies a multi-measure framework to understand variation in cognition and welfare ranges, i.e. how well or badly animals can fare, across farmed taxa. Every year, billions of animals worldwide are subject to farming practices that impact welfare, such as tail docking in pigs, beak trimming in chicken, and fin clipping in fish (Allen & Perry, 1975; Sutherland et al., 2008; Uglem et al., 2020; Franks et al., 2021). Given limited resources and the complexity of the challenge, societal efforts and decisions around funding allocations to reduce animal suffering are largely ad-hoc, lacking the rigorous and systematic approach critical to maximising welfare improvements. Insofar as the decision requires comparing welfare impacts across taxa, decision-makers need tools that are not currently available, including a framework for modelling differences in pain, suffering, and related cognition. However, making robust interspecies comparisons about cognition and welfare is exceptionally complex, owing to often-contending ethical, methodological, and practical considerations. As this knowledge is scattered across a broad literature, a key starting point is a comprehensive synthesis across species. While within-species taxa reviews exist (e.g., Marino, 2017; Lambert et al., 2017), they are not sufficiently broad in scope to address the present challenge of comparing between taxa.

To remedy this, the present review takes an interdisciplinary approach across animal welfare, comparative psychology, veterinary science, and philosophy (with an author from each field) to provide a comprehensive multi-taxa and multi-measure review of the empirical evidence on cognition and welfare ranges. This work is particularly timely because such reviews can lead to critical changes in animal welfare legislation. For example, in 2020, the UK government commissioned a report highlighting the compelling evidence for sentience in cephalopods and decapods (Birch et al., 2021, Sidebar 1), which led to the Animal Welfare (Sentience) Bill being extended to include both invertebrate groups. Here, we evaluate cognition and welfare ranges across 10 of the most extensively farmed animals: pigs, chickens, carp, salmon, octopus, shrimp, crabs, crayfish, bees, and silkworms. To investigate interspecies variation, we defined and assessed 90 empirically detectable proxies of cognition and welfare ranges (henceforth ‘proxies’) relating to behavioural flexibility, cognitive sophistication, and general learning. The result of this literature assessment was a comprehensive table of ratings based on references relevant, where available, for each proxy and taxa, giving an overall output of >1000 references (details in section 3.2). Fewer than 20 of 90 proxies identified had been tested across the majority of the 10 taxa, so we refined our review into a subset of these proxies in two overlapping catalogues (A and B). Catalogue A contains 10 ideal proxies, providing an optimal suite of proxies most closely linked to welfare. However, as the empirical data for these proxies were lacking for many taxa this negated interspecies comparisons. We thus created Catalogue B, containing a less-optimal suite of 10 proxies, some with weaker links to welfare. The Catalogue B proxies were selected as those with sufficient empirical data to enable interspecies comparisons.

### Sidebar 1: Example of interdisciplinary animal welfare review with legislation impacts - Birch et al. (2021)

The UK government recently commissioned a team of scientists to review the evidence of sentience – the capacity to experience emotions, with a focus on pain – in two invertebrate groups, cephalopods and decapods. The scientists used eight interdisciplinary criteria for determining sentience. The first four focused on whether the animal’s nervous system could support sentience. Specifically, whether the groups could (i) detect harmful stimuli; (ii) transmit those signals to the brain; (iii) process the signals in integrative brain regions; and (iv) change the nervous system’s response when exposed to painkillers or anaesthetics. The four remaining criteria focused on behaviour and cognition. Specifically, whether the groups could (v) trade-off risks of injury against opportunities for reward; (vi) tend to specific sites of injury using self-protective behaviours; (vii) learn to avoid harmful stimuli; and (viii) learn to value painkillers or anaesthetics when injured. After reviewing over 300 studies, the researchers found strong and diverse evidence of sentience in both groups. For example, exposure to acid caused crabs and octopuses to scratch and shield the affected area but self-protective behaviours ceased when exposed to an anaesthetic; crayfish exposed to repeated electrical fields showed anxiety-like mental states; and injured octopuses learned to favour locations where they could self-administer an anaesthetic. Results from this review led to including both cephalopods and decapods in the UK’s Animal Welfare (Sentience) Act 2022. Other countries including Norway, Sweden, and New Zealand have already given invertebrates legal protection, but many countries remain to recognise invertebrates as sentient.

Our review adapted the rating methods developed by Rethink Priorities (2020) and Birch et al. (2021) to compare the strength of the evidence against each proxy. For each taxa, we thus constructed a cognition and welfare range profile, with an overall judgement score based on current evidence, and discussed the implications of comparisons across the taxa. This process identified which proxies are supported by quantitative evidence and which should be prioritised for future research and funding. From this analysis, we proposed future research experiments for specific proxies to enable informative comparative research, including cognitive bias and inhibitory control related paradigms. We believe this approach will assist researchers, including potentially big-team science collaborations (Sidebar 2), to efficiently target existing knowledge gaps. We hope it will help streamline funding allocations and other key decisions that are required to improve the welfare of farmed animals.

### Sidebar 2: Big-team Science (BTS) Collaborations

There is a recent drive for the development of large collaborative, international networks aiming to promote Open Science practices, pool resources, and enable greater cross-species comparisons, with larger sample sizes and species representations to help remedy potential reproducibility and generalisation issues (Coles et al., 2022; Lambert et al., 2022). Examples of big-team science (BTS) projects focusing on non-human species include: <u>ManyPrimates, ManyBirds, ManyDogs, ManyGoats</u> and ManyFishes. These projects vary in terms of their study focus, however, efforts are being made to devise ‘ManyMany’ studies, where BTS projects plan to combine efforts in collaboration with <u>ManyBabies</u> and <u>Psychological Science Accelerator</u> to facilitate comparisons across humans, non-human primates, birds, fish, and others (Coles et al., 2022). Some current topics include: ManyPrimates on working memory (ManyPrimates et al. 2021), delay of gratification and inference by exclusion; ManyBirds on neophobia (responses to novelty; Miller et al., 2022); ManyGoats on responses to human attentional states; ManyDogs on dog-human social communication. These studies result in huge samples and greater statistical power compared to smaller studies that traditionally include several researchers and a single lab. For example, 400+ subjects across 40+ species and 30+ sites in ManyPrimates et al. (2021). Future studies may be driven by project core teams or collaborators. There is potential and scope to consider future studies of relevance to applied welfare and conservation, such as proxies highlighted in the current review, either through existing BTS projects, cross-BTS studies or development of a new BTS projects focusing on welfare, for instance, across farmed taxa.

## 2. COGNITION AND WELFARE

### 2.1 What is animal welfare and cognition?

Welfare may be defined as an animal’s state while responding to environmental challenges (Broom, 1996). There are various theories of animal welfare, but here we focus on hedonism, i.e., welfare determined by positively and negatively valanced experiences (Bruckner, 2020). In order to assess an animal’s welfare, it is necessary to make an objective assessment of a subjective state (Sandøe & Jensen, 2012), largely requiring reliance on measurable proxies for welfare. It is broadly agreed that no single proxy measure is sufficient for determining welfare (Botreau et al., 2009; Mellor & Beausoleil, 2015). As such, we integrate a variety of different proxies to create a welfare range profile (i.e., how well or badly an animal can fare) for each of our farmed taxa. Whether welfare ranges vary across taxa intersects with the theory that different species vary in their capacity to experience emotions. This concept has been referred to as the ‘emotional capacities claim’, which implies that animals with stronger emotional capacities possess larger welfare ranges (Višak 2017). In a similar vein, welfare often refers to both physical and mental needs, for example, the extent of awareness of an internal state when in pain, may determine how much the animal is actually suffering (Duncan & Petherick, 1991). This definition highlights the need to consider cognition, which can be broadly defined as including perception, learning, decision-making, and memory (Shettleworth, 2010). Cognition, therefore, includes the animal’s perspective when assessing welfare (Ferreira et al., 2021). For this reason, we also include cognitive assays to create our welfare range profiles. Cognitive assays of particular relevance to welfare include learning ability, preference tests, memory workload, capacity to recollect memories, behaviours associated with noxious stimuli, and cognitive bias (Brydges & Braithwaite, 2008). Some examples of research linking cognitive measures with welfare implications in chickens are included in Table 1; however, it is important to note that there is a general lack of fundamental, cognitive research addressing welfare issues (Fijn et al., 2020).

**Table 1.** Examples linking cognition and welfare in chickens.

| Proxy | Finding | Reference |
| --- | --- | --- |
| Individual differences | More fearful/ anxious chickens show reduced space use | Campbell et al., 2016 |
| Learning/ Individual differences | Low-ranging chickens show stronger food conditioning (place preference conditioning). Application: train to associate food or conspecifics with range | Ferreira et al., 2020 |
| Learning | Social learning of feather plucking in chickens. Application: increased use of visual barriers between plucking and non-plucking individuals, remove feather plucking individuals from flock, train 'demonstrator' birds that peck appropriate stimuli to encourage natural alternative | Zeltner et al., 2000; Freire, 2020 |
|  | behaviour to plucking |  |
| Learning/ Taste aversion | Spray feathers to reduce plucking in chickens. Application: alter taste/smell with sucrose vs quinine | Harlander-Matauschek et al., 2008 |
| Learning | Chronic hunger reduces learning in chickens (Y-maze) | Buckley et al., 2011 |
| Learning/ Navigation/ Individual differences | Adult hens that do not use outdoor areas were slower to learn the T-maze task compared to outdoor-preferring hens | Campbell et al., 2018 |
| Mental representation (of missing resources) | Rearing in a spatially complex environment leads to increased space use in chickens | Freire, 2020 |

### 2.2 What is a cognition and welfare range?

A cognition and welfare range (CWR) essentially refers to how well or badly animals can fare. It describes animals’ respective capacities for valanced experiences and can be used to assign relative moral weights to different species based on those capacities. For instance, one could understand a moral weight as the amount or range of welfare a species can realise, produce, or generate, from the best to worst welfare states possible. Therefore, CWR relates to how much welfare can be realised by individuals within a given species. This approach suggests that while every unit of realised welfare counts the same, some species may possess a larger number of possibly realised welfare units than others, i.e. a larger welfare range. In order to assess CWR, it is necessary to measure variation in capacities of relevance to welfare, for which the proxies provide some evidence. It assumes that animals with relatively large welfare ranges can be harmed to greater degrees (e.g., experience greater suffering) than animals with relatively small welfare ranges (outlined in Gaffney et al., 2022).

### 2.3 How to and why make interspecies comparisons

Resources for improving animal welfare in farming, laboratory, and other captive settings are often limited. As such, decisions need to be made around prioritisation of species and the focus of research and management protocols. These decisions are often made largely without rigorous evidence, as relevant tools are lacking. There remains a need for interspecies comparative tools that are grounded in empirical data, such as those relating to cognitive differences. For example, there has been a recent drive for the reduction of the use of non-human primates in invasive research as well as increased legal protection in several countries. This shift is partly due to increasing evidence for complex cognition and behaviour in these animals (Padrell et al., 2021). However, invasive work continues with many other taxa, such as rats and dogs, which arguably show similarly high levels of cognitive abilities, for instance, metacognition (rats: Foote & Crystal, 2007; dogs: Belger & Bräuer, 2018).

There are evident ethical, methodological, and practical considerations to be accounted for when making interspecies comparisons of cognition and welfare (Gaffney et al., 2022; Fischer, in press). One starting place is to utilise the field of comparative psychology, which investigates the evolution of cognition, for instance, involving comparisons of performance in cognitive tasks across different taxa (Chittka et al., 2012). Ideally, comparisons should utilise similar experimental paradigms and measures, while accounting or adapting for physical, social, and ecological differences between species. Large-scale cross-species comparisons of cognition are limited. Nevertheless, within the past decade there has been a drive to conduct big-team science collaborations with comprehensive comparisons of specific cognitive abilities across species. For example, short-term memory has been compared in 41 primate species (ManyPrimates et al., 2022), neophobia (responses to novelty) in 10 corvid species (Miller et al., 2022) and self-control (specifically inhibitory control) across 36 mammal and bird species (MacLean et al., 2014).

As it stands, many studies include single-species investigations or cross-species comparisons with a small number of taxa. Making comparisons between species where methodologies differ considerably is problematic. Nevertheless, providing limitations are acknowledged and conclusions are tentative, interspecies comparisons can be made on existing research through focused literature reviews, as we outline below.

## 3. METHODS

### 3.1 Species/ Taxa and Reviewers

We included a wide range of commonly farmed species and their families, namely: Suidae (inc pigs), Phasiandae (inc chickens), Cyprinidae (inc carp), Salmonidae (inc salmon), Octopodidae (inc octopus), Penaeidae (inc shrimps), Portunidae (inc crabs), Cambaridae (inc crayfish), Apidae (inc bees), and Bombycidae (inc silkworm). There were six reviewers (RM, MS, AKS, LPG, JML, AT), each responsible for 1-4 taxa, and all were experienced in differing areas of animal welfare and comparative cognition or had related research and practical experience. Literature reviews for all target taxa, except Octopodidae, were completed in June 2022, with Octopodidae completed in August 2022.

### 3.2 Literature Review

#### 3.2.1 Full Review: 90 Proxies

To investigate interspecies variation, we assessed 90 empirically detectable proxies of cognition and welfare ranges (henceforth proxies) relating to cognition, behaviour, anatomy, physiology, and welfare. We used a variation of Delphi method (Linstone & Turoff, 1975), a form of structured deliberation involving a panel of five experts (philosophers, comparative psychologists, neuroscientists), to select the full set of proxies. Each panel expert provided a list of proxies, which were then discussed regarding their merits and relevance, and the final lists were combined to create the full 90 proxy list in Dec 2021. For each combination of taxon and proxy, we reviewed the existing literature across 10 taxa to determine whether there was sufficient scientific data and, based on this, whether it was possible to assess the likelihood of whether a taxon possessed a proxy. We used: 1) Google Scholar (soft search) and 2) Web of Science (hard search), as well as recent taxa-specific reviews and cited references.

Taxa were listed at the family level. If, for a given proxy, the target family had not been studied, we expanded the search to similar families in the same order that had been studied. Similarly, if a given proxy had not been studied, though a similar proxy had been, we included the latter, though noted clearly if this occurred. An example of search terms for chickens included: *‘Phasianidae’*, ‘chicken’, ‘junglefowl’, ‘pheasant’, ‘bird’. An example of search terms for the proxy self-control included: ‘self-control’, ‘delay of gratification’, ‘inhibitory control’, ‘behavioural flexibility’, ‘reversal learning’. Fifteen of the 90 proxies (17%) within each taxa were reviewed independently by a second person (not involved with initial lit review for that taxon) and cross-checked to ensure inter-rater reliability.

#### 3.2.2 Reduced Review: ‘Catalogues A and B’

A large number of proxies had missing data for at least one taxon (fewer than 20 of 90 proxies had been tested across the majority of the 10 taxa) so we refined our review to a) focus on the most welfare relevant proxies and b) enable interspecies comparisons using existing research. We grouped a subset of the 90 proxies into two overlapping catalogues (A and B, Figure 1). Catalogue A contained 10 ideal proxies, providing an optimal suite most closely linked to welfare. The selection for Catalogue A was based on a combination of criteria typically used to assess consciousness (Birch et al., 2020) and sentience (Birch et al., 2021; Sidebar 1) in non-verbal animals. Such criteria are relevant because the question of interspecies comparisons is largely about the comparative level of consciousness or sentience of different animals, since we assume that animals with stronger emotional capacities have larger cognitive and welfare ranges (i.e., emotional capacities claim, Višak, 2017). Several cognitive measures that have been linked to emotional capacities include learning ability, preference tests, and cognitive bias (Brydges & Braithwaite, 2008). However, as the empirical data for these proxies were lacking for many taxa, this negated interspecies comparisons. We thus created Catalogue B, containing a less-optimal suite of 10 proxies, some with weaker links to welfare.

**Figure 1:**
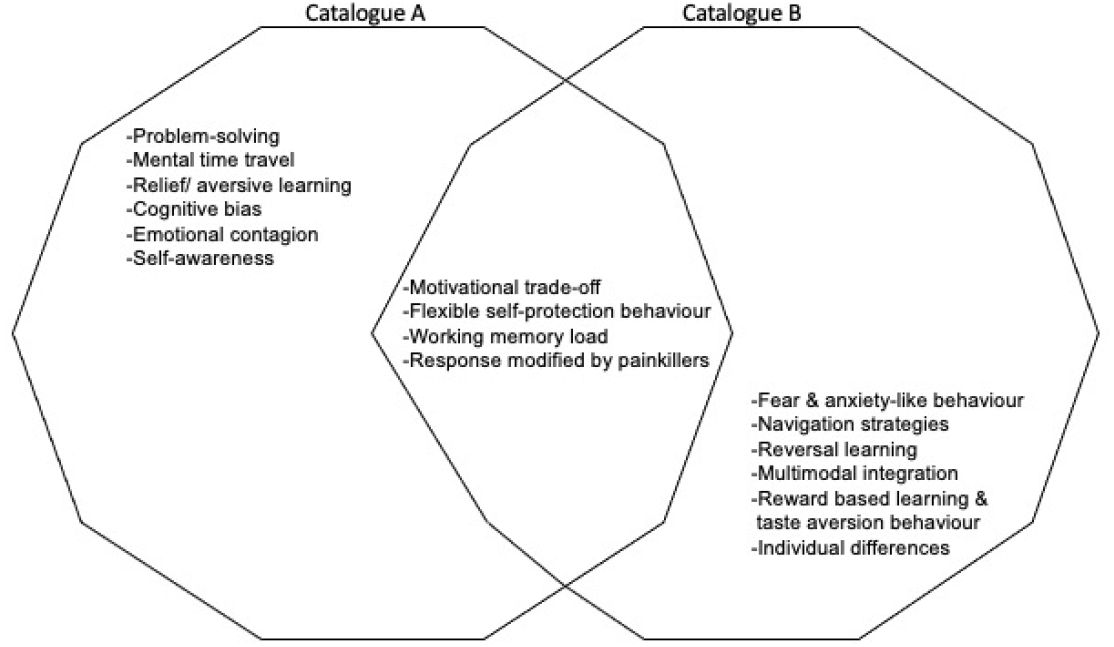
Catalogues A and B consisted of 16 proxies representing cognition and welfare range. Catalogue A provided an optimal framework, containing proxies most closely linked to welfare. Catalogue B provided a less-optimal framework, containing some proxies that are more weakly linked to welfare but are useful nonetheless as they facilitate direct interspecies comparisons via the availability of empirical data. Some proxies overlap both catalogues.

The Catalogue B proxies were selected as those with sufficient empirical data to enable intra- and interspecies comparisons. As such, we selected only those proxies that had been tested in the majority of the 10 taxa, regardless of rating, after filtering out those with many ‘unknown’ (i.e. insufficient evidence) ratings. The proxies in both catalogues tend to fall within the larger categories of behavioural flexibility, cognitive sophistication, and general learning. See Table 2 (glossary) for proxy definitions and Figure 1.

**Table 2.** Glossary of Catalogue A and B proxies.

| Catalogue | Category and Proxy | Definition |
| --- | --- | --- |
| A | Cognitive sophistication:<br>Mental time travel (incl. episodic-like memory, source memory, self-control, future planning, and memory integration) | Mental time travel is the ability to travel backwards and forwards in time in the mind's eye, to remember the past based on what happened, where and when (i.e., episodic-like memory, also consider source memory); and plan for the future (Tulving, 1983). Episodic-like memory is considered to be the precursor for future planning, as it functions as a memory database to predict future scenarios (Clayton et al., 2003; Schacter et al., 2012). Self-control, which is part of executive function, is an integral part of future planning because an individual must overcome immediate gratification to fulfil future needs (Schnell et al., 2021). |
| A | General learning:<br>Relief/ aversive learning | Relief learning involves the ability to associate that a specific stimulus results in relief from a negative state (e.g., pain) or results in the offset of a negative reinforcer. In a similar way, aversive learning involves the ability to avoid aversive stimuli that causes a negative state. There is a type of relief-memory that relates to something that happens after a painful event finishes, at the moment of the so-called relief. This relief can both increase the learning of the cues associated with the disappearance of the threat and reinforce those behaviours that helped to escape it (Gerber et al., 2014). |
| A | Cognitive sophistication:<br>Cognitive bias (incl. judgement, attention, memory) | Cognitive bias in animals is a pattern of deviation whereby inferences about new situations might be affected by irrelevant information or emotional states. There are several types of cognitive bias, which are measured using different methods. These include assessing whether individuals experiencing negative affect make more pessimistic judgements about ambiguous stimuli (i.e., judgement bias), pay more attention to negative stimuli (i.e., attention bias), and are more likely to remember negatively valenced memories (memory bias) than positively valenced memories (Eysenck et al., 1991; Wright & Bower, 1992; Paul et al., 2005). |
| A | Behavioural flexibility: Problem-solving | The ability to solve problems is important for animals to respond to a rapidly changing environment. Specifically, animals use problem-solving skills to avoid predators and obtain access to important food sources, shelter, or mates (Pérez Fraga et al., 2021). Problem-solving does not have to be a complicated process, and can rely on animals exploring their environment, learning and remembering information. However, problem-solving can also involve complex cognitive abilities such as logical reasoning, causal reasoning, and future planning (Andrews, 2021). Notice that different species, populations, or even individuals can solve problems in different ways. |
| A | Cognitive sophistication:<br>Emotional contagion | Emotional contagion occurs when an individual matches the emotional state with another individual or when emotions are transferred between individuals (Perez-Manrique & Gomila, 2022). It can be seen as evidence of more sophisticated emotions, and perhaps as a precursor to the capacity for |
|  |  | empathy, which indicates social cognition or even theory of mind (Reimert et al., 2013). |
| A | Cognitive sophistication: Self-awareness (incl. self-recognition i.e., MMR, knowledge of own mental states, knowledge of others' mental states i.e., theory of mind, body awareness, experience projection) | Self-awareness suggests an understanding or recognition of self. Self-recognition in non-human animals is generally measured via the mirror mark test (Gordon, 1970). One complementary avenue to measure self-recognition is to investigate body self-awareness. Body awareness is the ability to discriminate between body and non-self-stimuli, specifically understanding that an individual's body is distinctly different from the surroundings (Moore et al., 2007; Brownell et al., 2007; Dale & Plotnik, 2017). Self-awareness is often linked to other complex forms of cognition including empathy and perspective-taking. Perspective-taking, also termed theory of mind, is the ability to understand and consider another individual's mental state i.e., 'mind-reading' (Premack & Woodruff, 1978), including knowledge, desires and beliefs that motivate others' action (Krupenye, 2021). |
| A & B | Behavioural flexibility: Flexible self-protective behaviour | Flexible protective behaviour is a type of non-reflexive reaction to injury in which an injured animal attempts to guard, groom, or otherwise tend to the injured body part. Examples include limping, wound rubbing, wound licking, and wound guarding (Elwood, 2019). Protective behaviour, in our sense, must be carefully distinguished from reflexive reactions known (in humans) that operate subconsciously, such as grimacing, rapid withdrawal, postural adjustments, and some paralinguistic features of vocalisation (Sekhon et al., 2017). |
| A & B | Cognitive sophistication: Working memory load | Working memory load is a short-term memory system, describing a limited capacity to hold information temporarily. Working memory load is important for reasoning and guiding decision-making as well as executing behaviour (Miyake & Shah, 1997; Diamond, 2013). It is also responsible for following goals and keeping track of multiple goals, by integrating a variety of information sources (Miyake & Shah, 1997). |
| A & B | General learning: Response modified by painkillers | Painkillers such as local anaesthetics, analgesics (i.e., opioids), anxiolytics or anti-depressants modify an animal's response to noxious stimuli in a way that suggests that these compounds attenuate the experience of negative affective states (i.e., pain or distress). |
| A & B | Behavioural flexibility: Motivational trade-off | Motivational trade-off involves an animal having to flexibly trade-off between two competing motivations. Specifically, an animal must be motivated to avoid a noxious stimulus, and this motivation must be weighed (traded-off) against other motivations (e.g., thirst, hunger, the need for shelter) in a flexible decision-making process. Motivational trade-off has been demonstrated in hermit crabs who were offered an opportunity to hide from an electric shock in either their preferred *Littorina* shells or their non-preferred *Gibbula* shells, meaning the crabs had to trade shock avoidance against shell preference (Magee & Elwood, 2016). |
| B | Cognitive sophistication: Fear-like and anxiety-like behaviour | The experience of fear is associated with certain physiological and behavioural responses. Behavioural markings of fear include fleeing, hiding, freezing, and suspending unnecessary bodily functions. Physiological reactions to fear can include elevated heart rate, hyperventilation, increased muscle tension, constriction of blood vessels, nausea, and dizziness (Stankowich & Blumstein, 2005). Anxiety is related to but distinct from fear. |
|  |  | Anxiety is sometimes said to be the result of danger that is perceived to be unavoidable (Öhman, 2008) or situations in which the threat is ambiguous or unknown (Belzung & Philippot, 2007). |
| B | Behavioural flexibility: Individual differences/ 'personality' | Personality refers to individual differences in characteristic patterns of behaving. Temperaments and personalities are integrated behavioural phenotypes and stable traits that are consistent over time and across situations, which are broad and consistent dimensions of individuality (Budaev, 1997). |
| B | Cognitive sophistication:<br>Navigation strategies | Animals need to avoid environmental hazards and locate food, water, and mates. Many animals need to return to fixed sites (such as dens, hives, breeding grounds, food caches, watering holes, foraging sites, or nesting beaches) multiple times over the course of their lives. To do so requires navigational skills. Notice that navigation strategies can vary from simple associative processes and innate behaviours to complex cognitive mapping and memory recollection. Navigational skills are relevant for welfare ranges insofar as some strategies indicate advanced forms of cognition, which is correlated with increased welfare range. Importantly, the evolutionary function of consciousness is thought to produce an integrated and egocentric spatial model to guide an animal as it navigates a complex environment (Barron & Klein, 2016). |
| B | General learning:<br>Reversal learning | Reversal learning is the ability to change behaviour rapidly and flexibly in response to changing circumstances, which is an important tool for survival in a rapidly changing environment (Izquierdo et al., 2017). |
| B | Cognitive sophistication:<br>Multimodal integration | Multimodal integration is the ability to process different sensory modalities (e.g., light, touch, smell, sound) and integrate them via the nervous system (New, 2002). |
| B | General learning:<br>Reward based learning and taste aversion behaviour | Learning based on reward is one type of associative learning, which involves learning about a relationship between two separate stimuli. Associative learning can be differentiated into classical and operant conditioning (Pearce & Bouton, 2001). Taste aversion learning is a form of associative learning, which involves a learned pattern of aversion to a specific food after it has been paired with an aversive stimuli (e.g., radiation exposure, injection of some toxic drug such as lithium chloride, exposure to a high intensity magnet, etc.) (Bernstein, 1999). Specifically, the animal associates a transient state of illness to the taste, odour, or other characteristic of a specific food item, which ultimately results in a long-term change in its perception of palatability (Snijders et al., 2021). |

### 3.3 Evidence Rating

Our review adapted the rating methods developed by Rethink Priorities (2020) and Birch et al. (2021) to compare the strength of the evidence and probability against each proxy. In our reduced review, each proxy was rated per taxa using two approaches, whereas in the full review, each proxy was ranked using the probability rating scale per Rethink Priorities (2020). The rating systems lend to similar outputs, though we used both approaches for continuity and to enable comparison with previous studies:

1. Rethink Priorities (2020) use a probability rating scale with 5 grades to assess whether the evidence suggests a taxon possesses a proxy: ‘likely no’ (0–25% credences), ‘lean no’ (>25–<50%), ‘unknown’ (50%), ‘lean yes’ (>50–<75%) and ‘likely yes’ (>75–100%). Note that these 5 ‘credence assessments’ represent the evaluation of whether a taxon displays or fails to demonstrate the proxy rather than the *extent* to which the animals possessed the proxy. ‘Unknown’ was the default assessment in cases where insufficient evidence was found for a particular taxon/proxy combination.
2. Birch and colleagues (2021) use a level of confidence to grade the quantity, reliability and quality of the available evidence. There are six confidence levels in this rating method: (i) very high confidence, (ii) high confidence, (iii) medium confidence, (iv) low confidence, (v) very low confidence, and (vi) no confidence. The ‘very high’ confidence rating illustrates that the weight of scientific evidence leaves no room for reasonable doubt that the proxy is present or absent. The ‘high’ confidence rating illustrates that the animals convincingly display or fail to demonstrate the proxy but there is some room for reasonable doubt. The ‘medium’ confidence rating illustrates that there are some concerns about the reliability of the evidence. The ‘low’ confidence rating illustrates that there is little evidence that the animals display or fail to demonstrate the proxy. Finally, the ‘very low’ and ‘no’ confidence ratings illustrate that the evidence is either considerably inadequate or non-existent, respectively.

We calculated an overall judgement score per taxa and across proxies, using a comparable approach to Birch et al. (2021), comprising the total percentage of ‘very high’ and ‘high’ ratings. The grading scheme allocates ‘very high’ or ‘high’ confidence that a taxon satisfies 87.5% of proxies as **very strong**, 62.5% as **strong**, 38.5% as **substantial** and < 38.5 % as **weak** evidence for a larger cognition and welfare range, indicating a greater capacity to experience enhanced negative (e.g., greater suffering) and positive (e.g., greater enjoyment) emotions. Note that Birch et al. (2021) focus on evidence for sentience, specifically pain, and proxies of particular relevance for this, with overall judgements determined by percentage of criteria met per taxa, whereas we focus on proxies relevant to cognition and welfare rather than sentience (See Sidebar 1 for more detail).

## 4. RESULTS

### 4.1 Full Review Output: 90 proxies

The final table comprised 90 proxies across 10 taxa (Fischer, 2022; Table S1 and S2). We note that, across these 90 proxies, there were only 7 ‘likely no’ and 8 ‘lean no’ ratings - most of which were for proxies that had not been tested in many taxa (other than parental care), therefore this was not a useful alternative means of reducing the table, i.e., based on range of rating.

### 4.2 Reduced Review: Catalogue A and B

We outline Catalogue A (ideal proxies, with limited empirical data) and B (less-optimal proxies, with empirical data) output and ratings in Table 3, Figure 2 (Catalogue A and B combined) and Figure 3 (a: Catalogue A & B combined, Catalogue B alone; b: Categories - behavioural flexibility, cognitive sophistication, general learning). Please note, we provide only 1 citation per taxa and proxy in Table 3 as an example as the primary goal was to establish trait presence and also due to the sheer number of references generated by the review (123 references with only 1 reference example). The full 90-proxy review output identified >1000 references.

**Figure 2.**
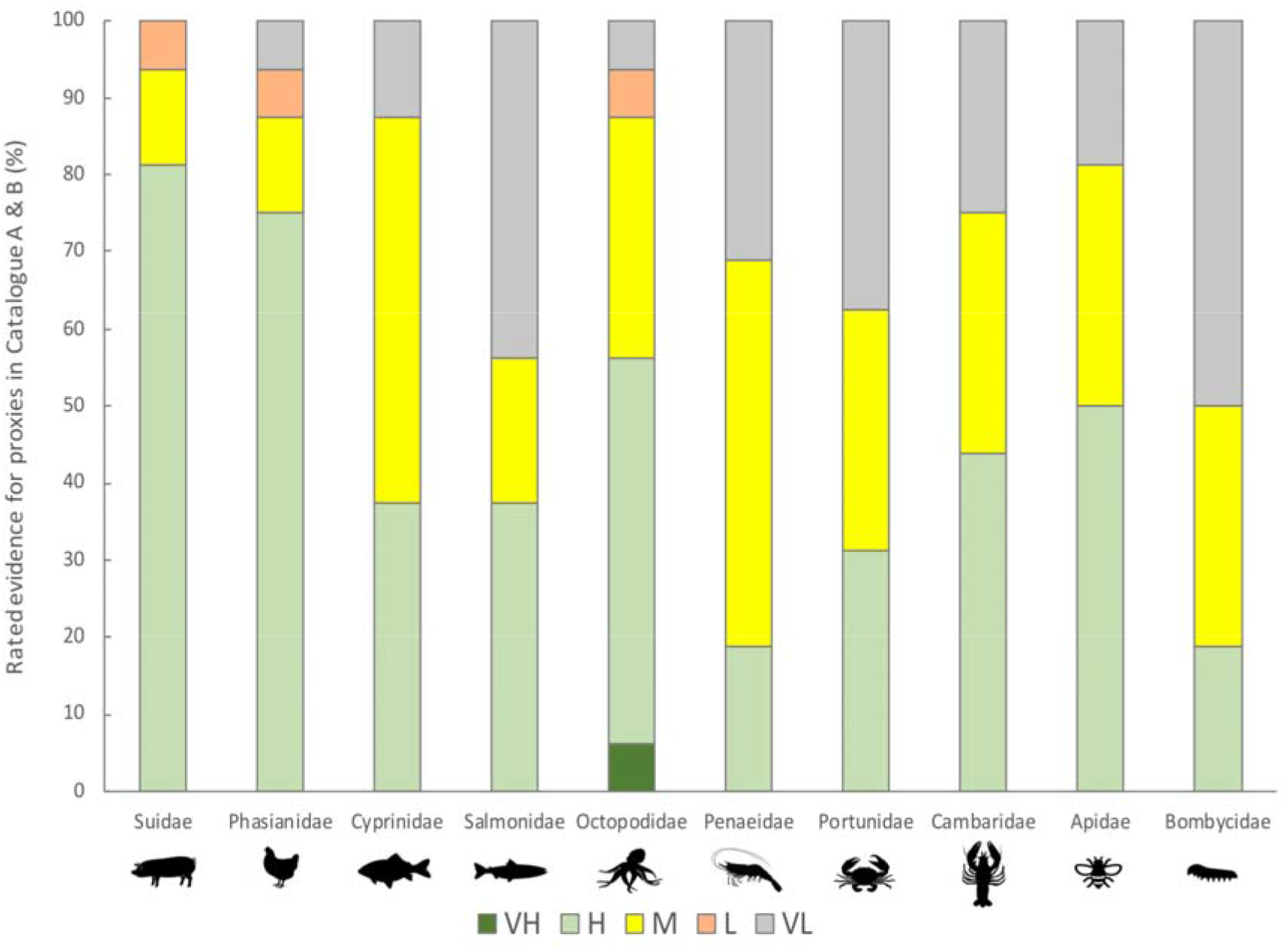
Rated evidence for proxies* listed in Catalogue A and B per taxa. Code: VH/ Very high: Large amount high-quality, reliable evidence; H/ High: Convinced based on evidence, with room for reasonable doubt; M/ Medium: Some concerns about reliability of evidence; L/Low: Little evidence; VL/ Very low: Seriously inadequate or non-existent evidence (per Birch et al., 2021 rating). Rating translates to: VH = yes, H = likely yes, M = lean yes, L = lean no, VL = unknown (per Rethink Priorities 2020 rating). * Table 3 lists the specific proxies.

**Figure 3:**
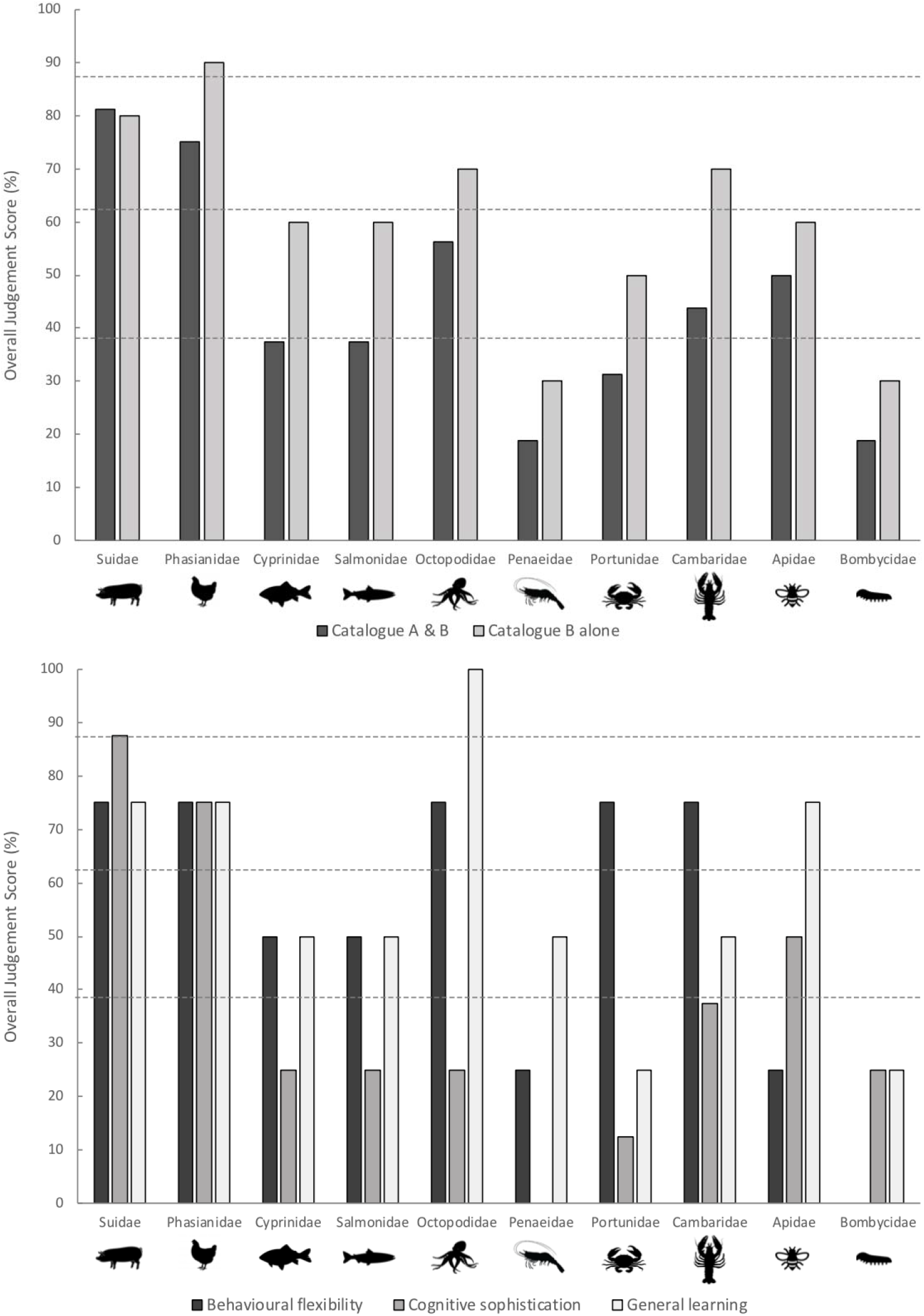
Percentage overall judgement scores (rated as high-very–high confidence out of all ratings) per taxa arranged across (a) Catalogues (A & B combined; B alone)* (b) Categories (behavioural flexibility, cognitive sophistication, general learning). Horizontal dotted lines reflect very high–high confidence in strength of evidence in descending order: at least 87.5% = Very strong; at least 62.5% = Strong; at least 38.5% = Substantial; values below the lowest horizontal dotted line < 38.5 % = weak (Birch et al., 2021).* Catalogue A alone is not represented in the graph because empirical data for these proxies were lacking for many taxa, thus it was not possible to make reliable interspecies comparisons with this catalogue alone at present.

**Table 3.**
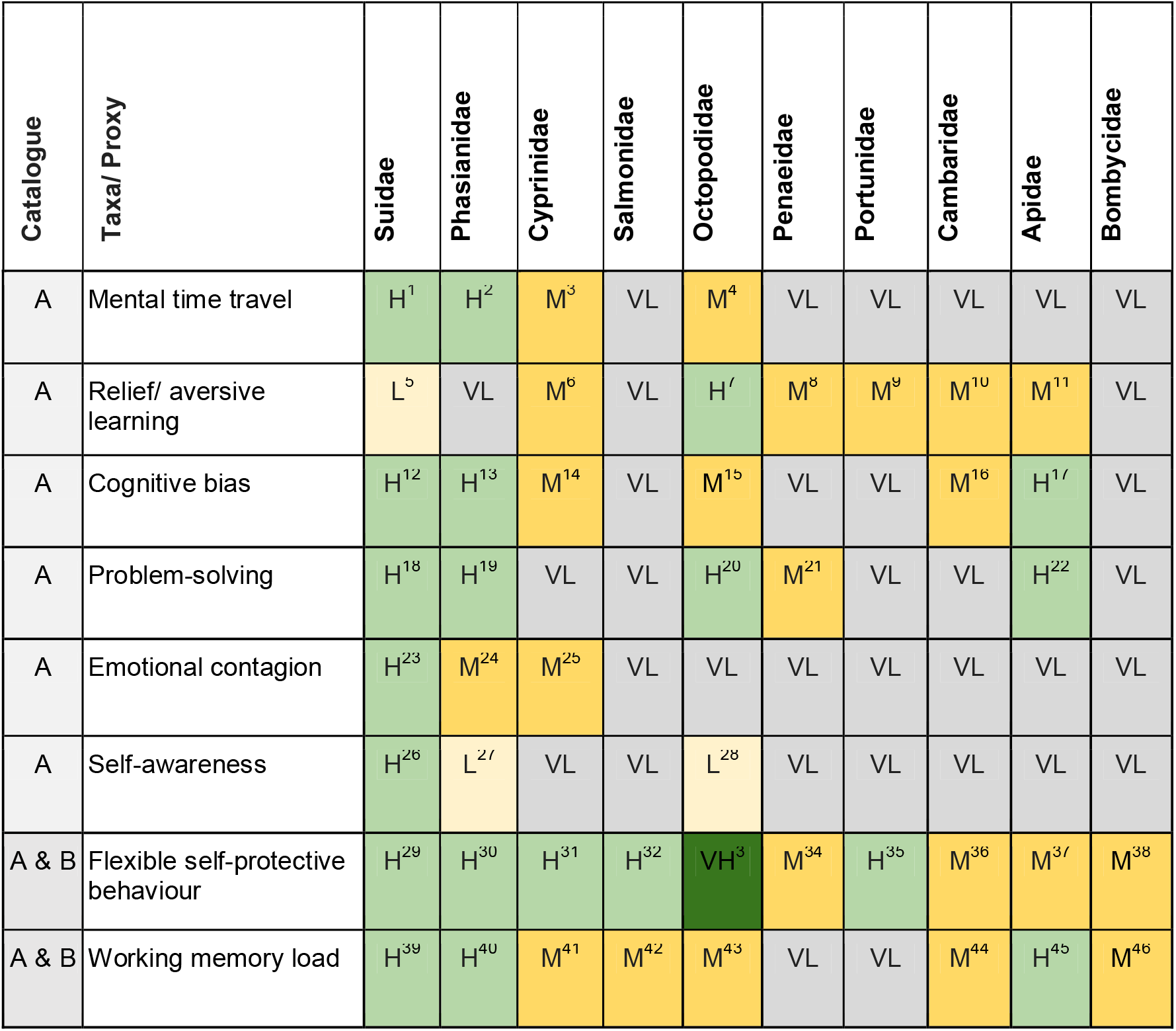
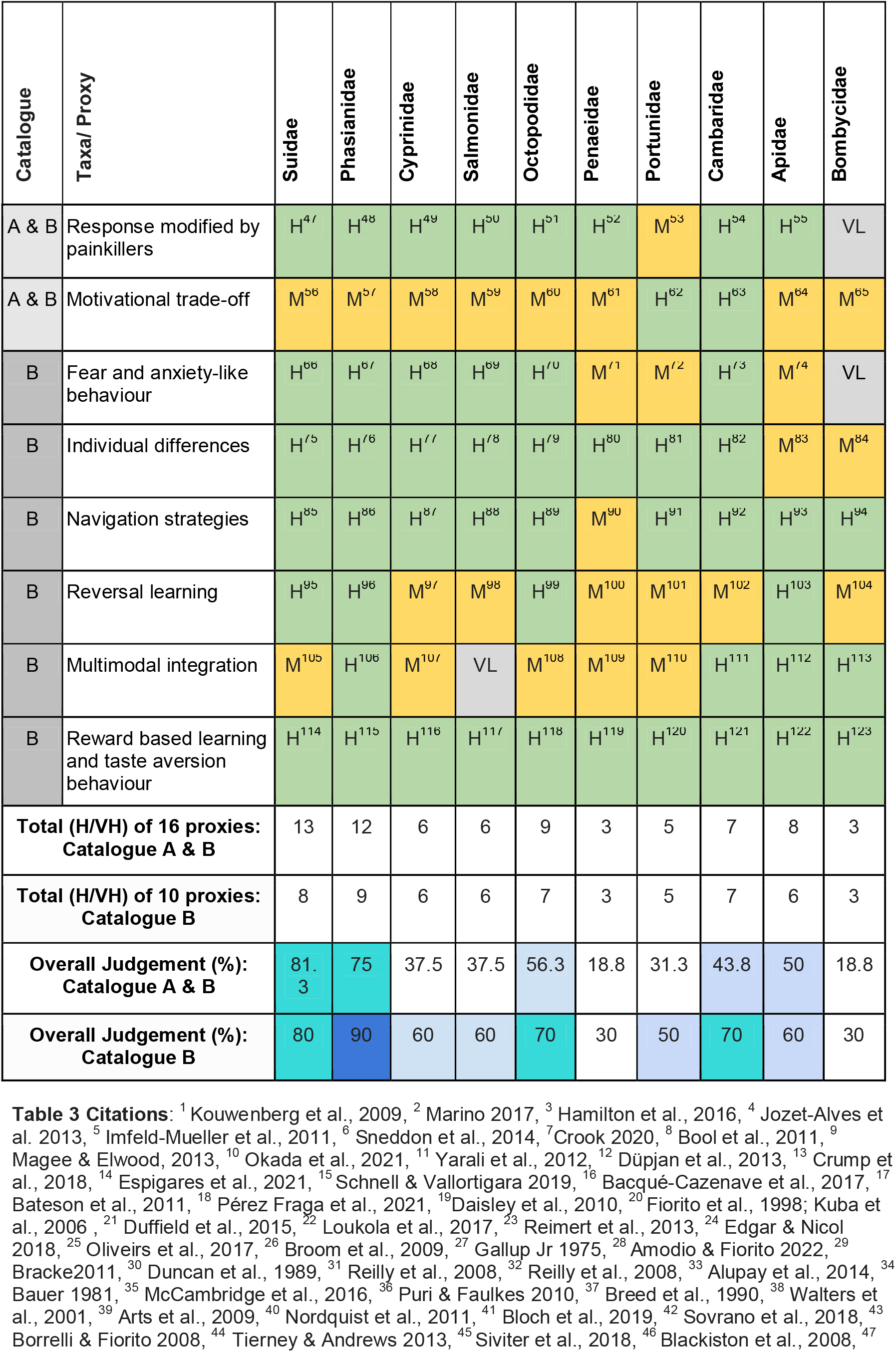
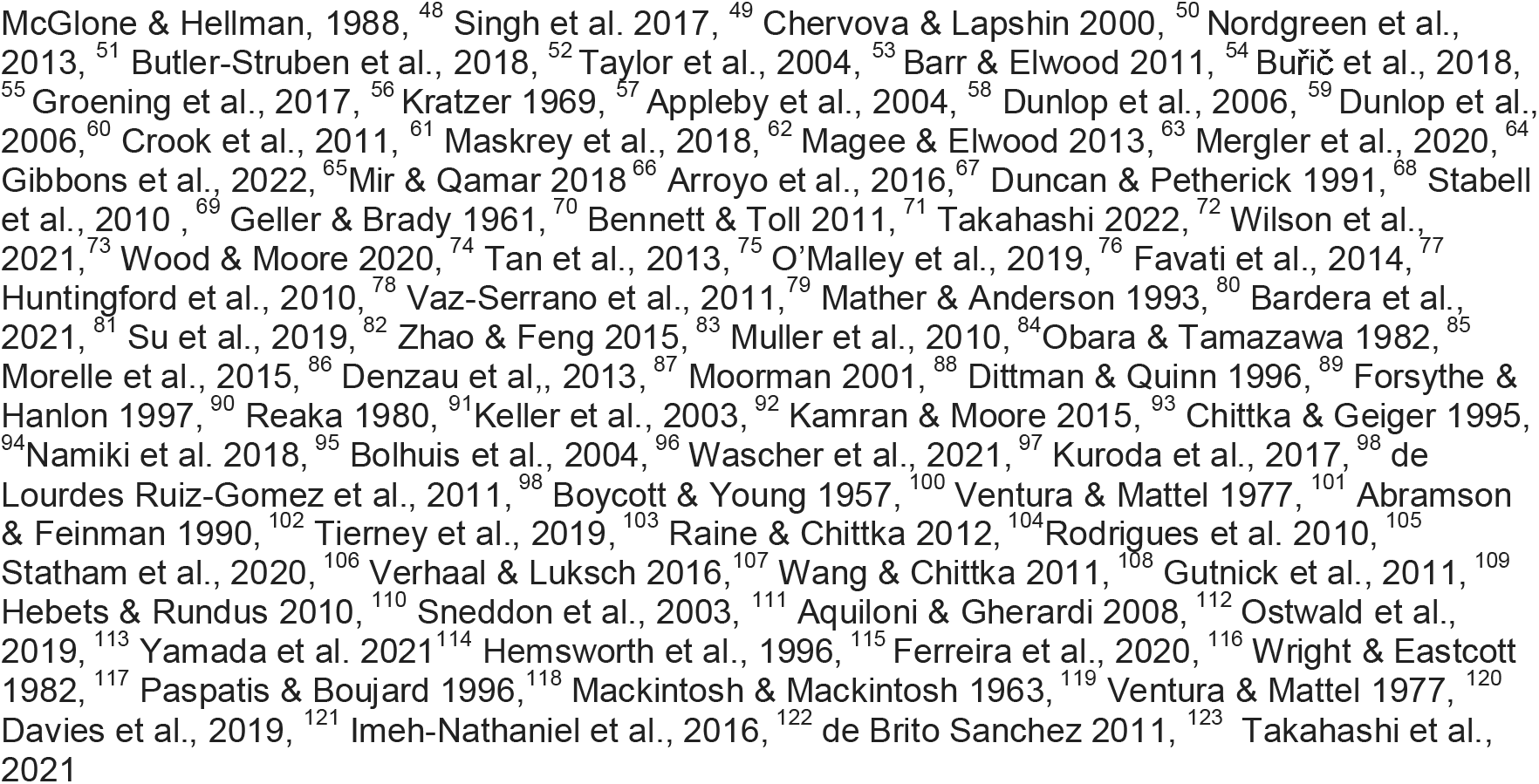
Cognition and welfare range proxies for Catalogue A and B per taxa. Code: VH/ Very high: Large amount high-quality, reliable evidence; H/ High: Convinced based on evidence, with room for reasonable doubt; M/ Medium: Some concerns about reliability of evidence; L/Low: Little evidence; VL/ Very low: Seriously inadequate or non-existent evidence (per Birch et al., 2021 rating). Rating translates to: VH = yes, H = likely yes, M = lean yes, L = lean no, VL = unknown (per Rethink Priorities 2020 rating). Overall judgement (% of VH & H out of overall ratings): 87.5% = Very strong; 62.5% = Strong; 38.5% = Substantial; < 38.5% = Weak (Birch et al., 2021).

### 4.3 Overall Judgement Score per Taxa

For Catalogue A & B combined (16 proxies), five of ten taxa meet criteria for either ‘strong’ or ‘substantial’ evidence for a larger cognition and welfare range, whereas for Catalogue B alone, eight taxa meet these criteria, one of which scored as ‘very strong’ (Table 3; Figure 3a). Specifically, for Catalogue A & B, *Suidae* (inc pigs) and *Phasianidae* (inc chickens) had a ‘strong’ overall judgement score, while *Octopodiae* (inc octopus), *Cambaridae* (inc crayfish), and *Apidae* (inc bees) had a ‘substantial’ score, and *Cyrinidae* (inc carp), *Salmonidae* (inc salmon), *Penaeidae* (inc shrimp), *Portunidae* (inc crabs), and *Bombycidae* (inc silk moths) only attained a ‘weak’ overall judgement score. For Catalogue B (10 proxies), *Phasianidae* had a ‘very strong’ score, *Suidae, Octopodiae, Cambaridae* had a ‘strong’ score, *Cyprinidae, Salmonidae, Portunidae*, and *Apidae* had a ‘substantial’ score, and *Penaeidae* and *Bombycidae* had a ‘weak score’.

## 5. DISCUSSION

### 5.1 Interspecies Comparisons: Similarities and Differences

The overall number of taxa with ‘strong’ to ‘substantial’ evidence supporting larger cognition and welfare ranges differed across our catalogues. We found stronger evidence for larger cognition and welfare ranges across taxa in Catalogue B alone, compared to Catalogue A & B combined. However, this is invariably because of a lack of research that focuses on proxies listed in Catalogue A, rather than clear evidence that some of the Catalogue A proxies are absent in our taxa. Given the lack of research in some taxa across the Catalogue A proxies, we propose that the Catalogue B is a more reliable catalogue at present. Within our target vertebrate taxa, there is ‘very strong’ to ‘strong’ evidence for larger cognition and welfare ranges in chickens and pigs, respectively. There is somewhat less evidence concerning carp and salmon, with the evidence in Catalogue B alone graded as ‘substantial’ and the evidence for Catalogue A & B combined graded as ‘weak’. Some of our target invertebrate taxa scored similarly to some of our higher scoring vertebrates in their cognition and welfare ranges. Indeed, for Catalogue B, pigs, octopus, and crayfish attained a ‘strong’ overall judgement score and chickens attained a ‘very strong’ overall judgement score.

Overall judgement scores for Catalogue A & B (Table 3) were more variable across the aforementioned taxa: pigs (81.3 %), chickens (75 %), octopus (56.3 %), and crayfish (43.8 %); but again, this is invariably because of a lack of positive evidence, rather than because of clear evidence that some of our taxa do not possess specific proxies in Catalogue A & B. Comparable scores, at least within Catalogue B, also exist between our target fish species (i.e., carp, 60 %; and salmon, 60 %), crabs (50 %), and bees (60 %). These findings add to existing evidence supporting recommendations for the audit and amendment of current housing, treatment, and other management decisions across *all* farmed animals, particularly fish and invertebrates, which tend to be amongst the least protected under legislation (e.g., Brown, 2014; Chittka, 2022).

We can assume that humans generally score the maximum possible judgement score on each proxy selected (i.e., 100%) as proposed by Rethink Priorities (2020). This would correspond with ‘very strong’ evidence under our rating system for a larger cognition and welfare range. In comparison, we also find ‘very strong’ evidence in chickens, at least for Category B proxies where more empirical evidence exists. These findings imply that at least some basic proxies measuring how well or badly a species may fare are comparable to some degree between humans and other species. Moreover, our data suggests that the same pattern exists between some of our vertebrate and invertebrate taxa (e.g., pigs, octopus, and crayfish). Attaining a similar cognitive and welfare range implies that these animals have comparable capacities for a wide range of valanced experiences, from negative (e.g., greater suffering) to positive (e.g., greater enjoyment).

### 5.2 Potential Implications and Recommendations

There are several implications and tentative recommendations from these findings with regard to prioritising funding and future research to improve welfare, as outlined in Figure 4. We highlight ‘key proxies’ in Catalogue A and elaborate on these in the ‘future research’ section.

**Figure 4:**
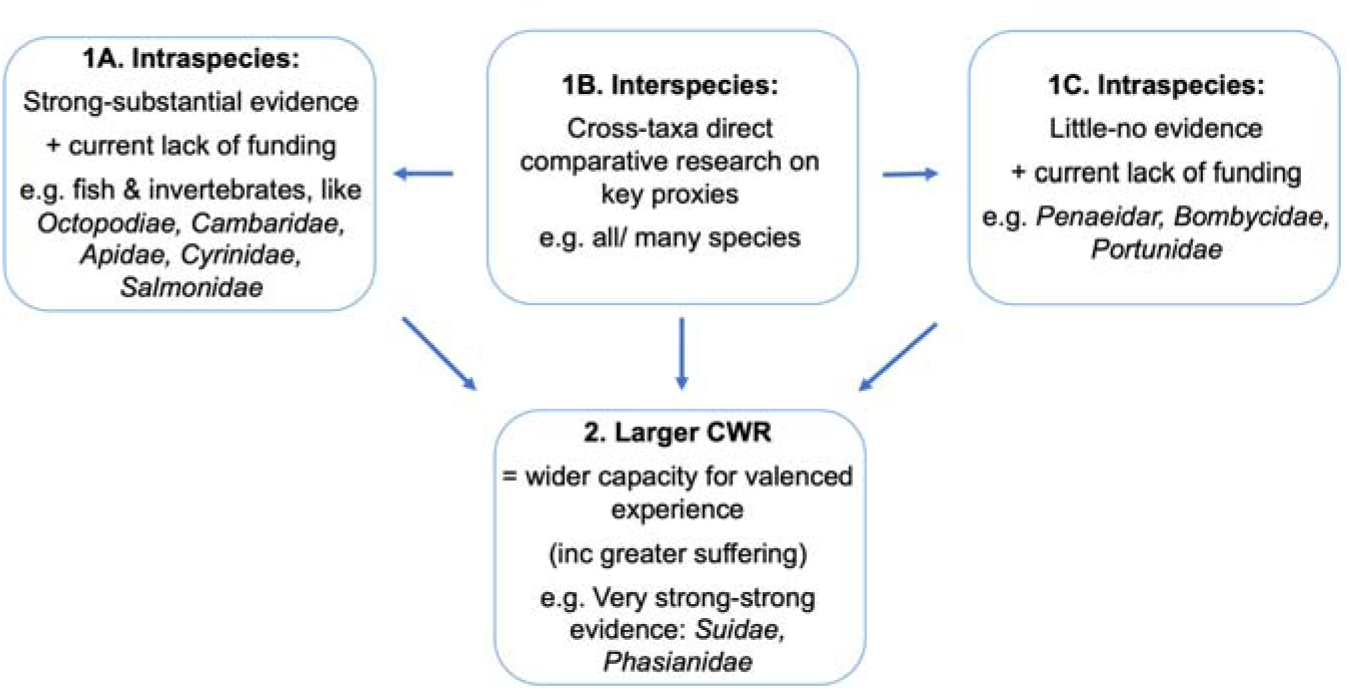
Recommendations for prioritising future funding and research.

Recommendations 1A) to 1C) should be prioritised first to inform generally as well as on recommendation 2) i.e., prioritising species with larger cognitive and welfare ranges (CWR). Recommendation 1B (interspecies comparisons) can inform on 1A, 1C and 2, though multi-species comparisons may require simplified experimental paradigms and take longer to complete given that large-scale multi-species studies typically require multiple collaborators and sites, compared to single-species studies. Therefore, 1A and 1C are also vital steppingstones to facilitate in-depth interspecies studies. The decision to prioritise species with some evidence (‘strong’ – ‘substantial’) compared with little to no evidence (‘weak’) of CWR may be further informed by current levels of funding allocations and other existing empirical data. For example, there is currently relatively little evidence for Portunidae (including crabs) to demonstrate a larger CWR.

These recommendations are not mutually exclusive. We advocate prioritising all farmed species to improve welfare. However, we also recognise that with limited financial support, funding and research allocation decisions have to be made and where possible, would benefit from being empirically based.

### 5.3 Limitations and Considerations

We highlight several potential limitations of our review; some of which are specific to the review, and others are more generally widespread across animal-related research though may impact the review findings. Within this review, one issue was the selection of suitable proxies for cognition and welfare range, which differentially influenced the overall judgement scores. For instance, overall judgement scores differed across catalogues for some taxa, i.e. *Phasianidae, Cyprinidae, Salmonidae, Octopodidae, Portunidae* and *Cambaridae* all obtained higher scores in Catalogue B alone compared to Catalogue A & B combined. Given that the scores are sensitive to proxy selection, our judgement scores should be treated with caution. It might be worthwhile placing more weight on the Catalogue B proxies, given that this is where more research has been conducted. However, we do not recommend dismissing the proxies listed in Catalogue A. It is widely suggested that evidence of psychological abilities such as mental time travel, problem-solving, and theory of mind suggests the animal possesses complex cognition (Emery & Clayton, 2004) and indicates a presence of sentience (Proctor, 2012). As such, these proxies are likely to be important for estimating CWR.

We took a two-pronged approach to ensure that we recorded a widespread search of the literature to gain the best possible overview of the current evidence for each proxy and taxa. However, it is possible that our literature reviews have not captured every relevant, existing study. Many studies, even within proxy, used different measures, designs and outputs, and thus, it was not possible to compare data directly. We were therefore required to make an informed judgement to provide a rating, with a subset of ratings cross-checked across two observers. We recognise that the current output (Table 3) will need to be updated as new studies are published or updated, however, we provide a necessary and comprehensive starting place (and potential methodologies) within this review article. One possible downside of the Birch et al. (2021) strength of evidence rating system is that it leans towards favouring presence, without allowing for indication of doubt in whether evidence for a proxy exists, as included in the Rethink Priorities (2020) rating system. For this reason, we used both rating systems and found similar outputs overall. Also, it is worth noting that our literature reviewers were aware of the study’s general theory and purpose (i.e. quantifying welfare ranges) when reviewing and rating proxies, and where possible, reviewed taxa that they were highly familiar with (e.g., experience conducting research and/or animal care). In the present study, interobserver reliability was conducted and confirmed; however going forward, we recommend that replication efforts conduct literature reviews while blind to study purpose.

Although some of the proxies have been tested in different species, they are rarely done so comparatively, which severely limits direct comparisons. For example, similar paradigms have been used to test for delay of gratification across species, but researchers often focus on differing measures and procedures that limit cross-species comparability (outlined in Miller et al., 2019 review of self-control in crows, parrots, and non-human primates). Efforts to expand on multi-species comparisons, for instance short-term/working memory in primates (ManyPrimates et al., 2019, Sidebar 2), may be a step towards remedying this in the future.

It is critical to keep in mind that ‘absence of evidence is not evidence of absence’ (Birch, 2017; Kuntsson & Munthe, 2017). Specifically, lack of evidence is likely to reflect lack of existing publications, rather than necessarily a confirmed lack of positive or negative evidence. The publication bias against negative findings leads to many studies that fail or show negative results not being published. Of the 90 proxies identified in the full table (Fischer, 2022; Table S1 and S2), fewer than 20 had been tested across the majority of the 10 taxa of focus. These areas highlight possible avenues for future empirical research and cross-species comparisons.

Similarly, as with many areas of science, the fields of animal cognition, behaviour, welfare science and others face some concerns and issues regarding replicability, low statistical power and sample sizes, as well as generalisation (Beran, 2018; Farrar et al., 2021; Open Science Collaboration, 2015). In response to this replication crisis, there is a push for greater use of Open Science practices (Munafò et al., 2017). This has also driven the development of big-team science projects (Coles et al., 2022; Sidebar 2). These issues impact on the reliability in interpreting some of the existing studies both within-species and between-species. For example, as outlined in Farrar et al. (2021) using inhibitory control (specifically the ‘cylinder’ task) as a case study.

We note that poor welfare can impact on cognition and behaviour, and thus it is possible that low cognitive performance in farmed animals may be a result of overall management practices, as opposed to being representative of typical species-level capacities. Similarly, study animals may be ‘STRANGE’ (e.g., their social background, rearing history) or ‘CRAMPED’ (e.g., have compromised health and development) hampering generalizability (Webster & Rutz, 2020; Cait et al., 2022). We remedied this to some extent by including a wider family focus (i.e., *Phasianidae*, as opposed to only chickens) to gather, where available, a wider representation of species and studies. Furthermore, at least for chickens, evidence is lacking for domestication leading to reduced cognitive or perceptual abilities in domestic chickens compared with their wild counterparts (i.e., red junglefowl) (Marino, 2017).

### 5.4 Future Research

Catalogues A & B represent outlines of proxies for prioritising in future (particularly comparative) research aimed at targeting welfare-relevant measures. In particular, cognitive bias, mental time travel (including self-control), relief/ aversive learning, emotional contagion, problem-solving, and self-awareness are all Catalogue A measures that are currently lacking in empirical data for most selected taxa (see Table 2 for proxy definitions). Furthermore, there are several Catalogue B measures that would benefit from direct comparative approaches, such as reversal learning and motivational trade-off, and expanding on memory related measures, such as episodic-like memory. For instance, episodic-like memories bring a past moment into the present, providing opportunity for individuals to recall details of these personal experiences. In this regard, negative memories can be especially powerful if an animal has the capacity to recollect the emotions that are linked to the experience, which is likely to lead to a larger welfare range.

We expand on two proxies, one per Catalogue, that may benefit from future research focus. Within cognitive bias (Catalogue A) tests, multiple study designs may be used, such as go/no-go, go/go (or active choice), or active choice with negative reinforcement methods (see reviews by Bethell et al., 2015 and Baciadonna & McElligott, 2015). Each method has various critiques, such as a large amount of training and confounding aspects of an animal’s internal state, like motivation or arousal, in go/no-go tasks. Go/go tasks may be more robust to such differences, though still require extensive training (Bethell et al., 2015). Tasks can also be adapted for cross-species comparisons by requiring different behavioural responses (e.g., nose poking, lever pressing, screen pecking) and varying sensory modalities (e.g., visual, auditory, textural cues) that are most appropriate for the study species. For instance, play related experiments have been tested across mammals, birds and insects, including recently using ball-rolling in bees (Dona et al., 2022). While modifications are necessary to support diverse taxa, it is important to design translatable tasks that facilitate cross-taxa comparisons. For instance, judgement bias tasks in humans commonly use secondary reinforcers (*e.g*., money - see Neville et al. 2021a) whereas judgement bias tasks in animals tend to use primary reinforcers (*e.g*., food - see Neville et al. 2020), making it challenging to compare results. Studies on humans should aim to use primary reinforcers (*e.g*., juice/salty tea) to make judgement bias tasks translatable between humans and animals (Neville et al. 2021b). By designing a series of translatable tasks, we can better draw conclusions about susceptibility to cognitive bias across taxa in the future.

Within reversal learning (Catalogue B) tests – a method for measuring inhibitory control – the methodologies often differ considerably despite being adapted for many species, making comparisons problematic. For example, the output measure may be learning speed, accuracy or error rates, or number of trials to reach criterion (acquisition and/or reversal phases). Similarly with cognitive bias tests, it can be easily modified to suit modality (e.g., colour or shape discriminations) and behavioural response. The complexity can also be increased, since some studies use additional reversals i.e., serial reversals tasks. Future research may look to standardise a methodology, including type of stimulus, criterions and output measures, and test across taxa. These outputs can then also be correlated with other factors, such as brain-to-body ratio and neuronal density (Olkowicz et al., 2016). For example, three North American corvid species differing in sociality were tested using a reversal learning colour discrimination task. Results revealed that highly social pinyon jays outperformed more solitary Clark’s nutcrackers and California scrub jays (Bond et al., 2007).

Further research should aim to directly correlate a) across different measures, such as cognition and welfare, and b) within related mechanisms, such as learning proxies such as social and contextual learning, or classical and operant conditioning. Cross-team collaborative approaches focusing on measures relevant and testable across a wide range of species will allow for more reliable inter-species comparisons of cognition, welfare, and other measures.

### Conclusion

Do certain animals have a greater capacity for suffering? This article presents and applies a multi-measure framework to understand variation in cognition and welfare range across 10 farmed taxa. For each taxon, we constructed a cognition and welfare range profile, with an overall judgement score, and discussed the implications of comparisons across the taxa. This process identified which proxies are supported by quantitative evidence and which should be prioritised for future research and funding. Our results reveal some variation in CWR across farmed taxa. Animals with larger cognitive and welfare ranges should be prioritised because we assume they have a greater capacity to experience enhanced negative (e.g., greater suffering) and positive (e.g., greater enjoyment) emotions. That being said, the research gaps are large, particularly in the proxies listed in Catalogue A; and thus, we are not yet in a position to construct comprehensive (i.e., data completeness) CWR profiles for all of our target taxa. Nevertheless, our analysis allows us to make broad, evidence-based comparisons with the data that is available. For example, evidence for proxies linked to behavioural flexibility is comparable across pigs, chickens, octopus, crabs, and crayfish (i.e., 75 %). Comparisons of this type can be made along all categories. Evidence for proxies linked to general learning is comparable across carp, salmon, shrimp, and crayfish (i.e., 50 %), whereas octopuses obtained a much stronger score in this category (i.e., 100 %). These conjectures may be overturned as researchers collect more detailed evidence, but it is a starting point.

Our analysis of current evidence also highlights gaps in the literature. To help bridge these gaps, we propose some future research experiments for specific proxies, which will enable informative comparative research. These include cognitive bias and inhibitory control related paradigms, both of which have been proposed as welfare-relevant measures. We believe this approach will assist researchers, including Big-Team Science collaborations (Sidebar 2), to efficiently target existing knowledge gaps and accelerate these objectives. Ultimately, this approach should help streamline funding allocations for welfare improvements and increase the overall welfare of farmed species.

## Acknowledgements

We would like to thank Richard Bruns, Marcus Davis, Adam Shriver, and Michael St. Jules for their discussion of the ideas that facilitated this approach to making interspecies welfare comparisons.

