## Supplementary Materials for "From Pigs to Silkworms: Cognition and Welfare across 10 Farmed Taxa"

**SUPPLEMENTAL MATERIALS**

**Table S1. Ratings: Full review with 90 proxies across 10 taxa per Rethink Priorities rating method**

| Proxies | **Pig** | **Chicken** | **Carp** | **Salmon** | **Octopus** | **Shrimp** | **Crab** | **Crayfish** | **Bee** | **Silkworm** |
| --- | --- | --- | --- | --- | --- | --- | --- | --- | --- | --- |
| A not B detour test | Likely yes | Likely yes | Unknown | Unknown | Unknown | Unknown | Unknown | Unknown | Likely yes | Unknown |
| Attention bias | Lean yes | Lean yes | Unknown | Unknown | Lean yes | Unknown | Unknown | Unknown | Unknown | Unknown |
| Body awareness | Lean yes | Unknown | Unknown | Unknown | Unknown | Unknown | Unknown | Unknown | Unknown | Unknown |
| Communication | Likely yes | Likely yes | Likely yes | Likely yes | Likely yes | Lean yes | Likely yes | Likely yes | Likely yes | Likely yes |
| Cooperative behaviour | Lean yes | Lean yes | Lean yes | Unknown | Lean yes | Lean yes | Lean yes | Likely yes | Lean yes | Lean yes |
| Cross-modal learning | Unknown | Likely yes | Lean yes | Unknown | Lean no | Unknown | Unknown | Unknown | Likely yes | Lean yes |
| Experience projection | Lean yes | Unknown | Unknown | Unknown | Unknown | Unknown | Unknown | Unknown | Unknown | Unknown |
| Gratification delay | Likely yes | Likely yes | Unknown | Unknown | Unknown | Unknown | Unknown | Unknown | Likely yes | Unknown |
| Individual differences / "personality" | Likely yes | Likely yes | Likely yes | Likely yes | Likely yes | Likely yes | Likely yes | Likely yes | Lean yes | Lean yes |
| Inhibitory control | Lean yes | Likely yes | Lean yes | Unknown | Lean yes | Unknown | Unknown | Unknown | Likely yes | Unknown |
| Judgment bias | Likely yes | Likely yes | Lean yes | Unknown | Unknown | Unknown | Unknown | Lean yes | Likely yes | Unknown |
| Memory bias | Unknown | Unknown | Unknown | Unknown | Unknown | Unknown | Unknown | Unknown | Unknown | Unknown |
| Mental Time Travel | Lean yes | Lean yes | Unknown | Unknown | Unknown | Unknown | Unknown | Unknown | Unknown | Unknown |
| Mirror mark recognition | Lean yes | Likely no | Unknown | Unknown | Lean no | Unknown | Unknown | Unknown | Unknown | Unknown |
| Motivational trade-off | Likely yes | Likely yes | Lean yes | Lean yes | Lean yes | Lean yes | Likely yes | Likely yes | Likely yes | Lean yes |
| Multimodal integration | Lean yes | Likely yes | Lean yes | Unknown | Likely yes | Lean yes | Lean yes | Likely yes | Likely yes | Likely yes |
| Navigation strategies | Likely yes | Likely yes | Likely yes | Likely yes | Likely yes | Lean yes | Likely yes | Likely yes | Likely yes | Likely yes |
| Numerical cognition | Unknown | Lean yes | Lean yes | Unknown | Unknown | Unknown | Unknown | Unknown | Likely yes | Unknown |
| Object permanence | Likely yes | Likely yes | Lean yes | Lean yes | Unknown | Unknown | Unknown | Unknown | Unknown | Unknown |
| Perspective taking | Lean yes | Unknown | Unknown | Unknown | Unknown | Unknown | Unknown | Unknown | Unknown | Unknown |
| Physical reasoning | Lean yes | Likely yes | Unknown | Unknown | Lean yes | Unknown | Unknown | Unknown | Unknown | Unknown |
| Problem-solving | Likely yes | Likely yes | Unknown | Unknown | Likely yes | Lean yes | Unknown | Unknown | Likely yes | Unknown |
| Response-slowing | Unknown | Unknown | Unknown | Unknown | Unknown | Unknown | Unknown | Unknown | Unknown | Unknown |
| Responses to novelty | Likely yes | Likely yes | Lean yes | Likely yes | Likely yes | Unknown | Unknown | Likely yes | Likely yes | Unknown |
| Reversal learning | Likely yes | Likely yes | Lean yes | Lean yes | Lean yes | Lean yes | Likely yes | Lean yes | Likely yes | Lean yes |
| Reverse reward test | Unknown | Unknown | Unknown | Unknown | Lean yes | Unknown | Unknown | Unknown | Unknown | Unknown |
| Shared intentionality | Unknown | Unknown | Unknown | Unknown | Unknown | Unknown | Unknown | Unknown | Unknown | Unknown |
| Social learning | Likely yes | Likely yes | Likely yes | Likely yes | Lean yes | Unknown | Unknown | Lean yes | Likely yes | Unknown |
| Socio-spatial cognition | Lean yes | Likely yes | Lean yes | Likely yes | Unknown | Unknown | Unknown | Unknown | Likely yes | Unknown |
| Symbolic representation of the world | Unknown | Lean yes | Unknown | Unknown | Unknown | Unknown | Unknown | Unknown | Unknown | Unknown |
| Temporal/ spatial discounting | Lean yes | Lean yes | Unknown | Unknown | Unknown | Unknown | Unknown | Unknown | Lean yes | Unknown |
| Theory of mind | Lean yes | Unknown | Unknown | Unknown | Unknown | Unknown | Unknown | Unknown | Unknown | Unknown |
| Tool use | Lean yes | Unknown | Unknown | Unknown | Likely yes | Unknown | Lean yes | Unknown | Lean yes | Unknown |
| Transitive inference | Lean yes | Likely yes | Unknown | Likely yes | Unknown | Unknown | Unknown | Unknown | Lean no | Unknown |
| Uncertainty monitoring | Unknown | Unknown | Unknown | Unknown | Unknown | Lean yes | Unknown | Lean yes | Lean yes | Unknown |
| Anxiety-like behaviour | Likely yes | Likely yes | Likely yes | Likely yes | Likely yes | Lean yes | Lean yes | Likely yes | Lean yes | Unknown |
| Associative learning from pain | Unknown | Likely yes | Likely yes | Likely yes | Likely yes | Unknown | Lean yes | Unknown | Lean yes | Unknown |
| Boredom-like behaviour | Likely yes | Likely yes | Unknown | Unknown | Lean yes | Unknown | Unknown | Unknown | Unknown | Unknown |
| Concept of death | Unknown | Unknown | Unknown | Unknown | Unknown | Unknown | Unknown | Unknown | Unknown | Unknown |
| Curiosity-like behaviour | Unknown | Unknown | Lean yes | Unknown | Lean yes | Unknown | Unknown | Unknown | Unknown | Unknown |
| Depression-like behaviour | Likely yes | Likely yes | Lean yes | Lean yes | Unknown | Unknown | Unknown | Unknown | Unknown | Unknown |
| Disgust-like behaviour | Lean yes | Likely yes | Unknown | Unknown | Unknown | Unknown | Unknown | Unknown | Unknown | Unknown |
| Displacement behaviour | Likely yes | Likely yes | Unknown | Unknown | Unknown | Lean yes | Unknown | Unknown | Lean yes | Unknown |
| Effects on exploratory behaviour from pain | Likely yes | Likely yes | Lean yes | Lean yes | Likely yes | Unknown | Unknown | Unknown | Unknown | Unknown |
| Emotional contagion | Likely yes | Lean yes | Lean yes | Unknown | Unknown | Unknown | Unknown | Unknown | Unknown | Unknown |
| Emotional reactions to learning | Lean yes | Lean yes | Unknown | Unknown | Lean yes | Unknown | Unknown | Unknown | Unknown | Unknown |
| Exhaustion-like behaviour | Likely yes | Likely yes | Likely yes | Likely yes | Likely yes | Likely yes | Unknown | Lean yes | Unknown | Lean yes |
| Fear-like behaviour | Likely yes | Likely yes | Likely yes | Likely yes | Likely yes | Lean yes | Lean yes | Likely yes | Lean yes | Unknown |
| Flexible self-protective behaviour | Likely yes | Likely yes | Likely yes | Likely yes | Likely yes | Lean yes | Likely yes | Lean yes | Lean yes | Lean yes |
| Friendship-like behaviour | Likely yes | Unknown | Lean yes | Unknown | Unknown | Unknown | Unknown | Unknown | Unknown | Unknown |
| Helping behaviour | Lean yes | Lean yes | Unknown | Unknown | Unknown | Unknown | Unknown | Lean yes | Lean yes | Unknown |
| Hyperalgesia | Lean yes | Lean yes | Unknown | Unknown | Likely yes | Unknown | Unknown | Unknown | Unknown | Likely yes |
| Jealousy-like behaviour | Unknown | Unknown | Unknown | Unknown | Unknown | Unknown | Unknown | Unknown | Unknown | Unknown |
| Joy-like behaviour | Unknown | Unknown | Lean yes | Unknown | Lean yes | Unknown | Unknown | Unknown | Unknown | Unknown |
| Liking-wanting dissociation | Unknown | Unknown | Unknown | Unknown | Unknown | Unknown | Unknown | Unknown | Unknown | Unknown |
| Loneliness-like behaviour | Likely yes | Likely yes | Lean yes | Lean yes | Unknown | Unknown | Unknown | Unknown | Unknown | Unknown |
| Love-like behaviour | Unknown | Unknown | Unknown | Unknown | Unknown | Unknown | Unknown | Unknown | Unknown | Unknown |
| Maternal response to offspring distress | Likely yes | Likely yes | Lean no | Lean no | Likely no | Lean no | Unknown | Unknown | Unknown | Likely no |
| Mourning-like behaviour | Unknown | Unknown | Unknown | Unknown | Unknown | Unknown | Unknown | Unknown | Unknown | Unknown |
| Panic-like behaviour | Likely yes | Likely yes | Lean yes | Likely yes | Lean yes | Unknown | Unknown | Unknown | Unknown | Unknown |
| Parental care | Likely yes | Likely yes | Lean no | Likely yes | Likely no | Likely no | Lean yes | Likely yes | Likely yes | Likely no |
| Play behaviour | Likely yes | Likely yes | Lean yes | Lean yes | Likely yes | Unknown | Unknown | Unknown | Lean yes | Unknown |
| Play vocalization | Likely yes | Unknown | Unknown | Unknown | Likely no | Unknown | Unknown | Unknown | Unknown | Unknown |
| Prioritizes pain response in relevant context | Likely yes | Likely yes | Lean yes | Likely yes | Lean yes | Likely yes | Lean yes | Unknown | Unknown | Unknown |
| PTSD-like behaviour | Lean yes | Unknown | Lean yes | Unknown | Unknown | Unknown | Unknown | Unknown | Unknown | Unknown |
| Relief learning | Likely yes | Unknown | Lean yes | Unknown | Likely yes | Lean yes | Lean yes | Lean yes | Lean yes | Unknown |
| Rescue behaviour | Lean yes | Lean yes | Unknown | Unknown | Unknown | Unknown | Unknown | Unknown | Unknown | Unknown |
| Response modified by painkillers | Likely yes | Likely yes | Likely yes | Likely yes | Likely yes | Likely yes | Lean yes | Likely yes | Likely yes | Unknown |
| Reward based learning | Likely yes | Likely yes | Likely yes | Likely yes | Likely yes | Lean yes | Likely yes | Likely yes | Likely yes | Likely yes |
| Sadness-like behaviour | Unknown | Unknown | Unknown | Unknown | Unknown | Unknown | Unknown | Unknown | Unknown | Unknown |
| Self-medication | Lean yes | Lean yes | Lean yes | Lean yes | Likely yes | Unknown | Unknown | Lean yes | Lean yes | Lean yes |
| Sensory-Affective Dissociation | Unknown | Lean yes | Unknown | Unknown | Likely yes | Unknown | Unknown | Unknown | Unknown | Unknown |
| Social buffering | Lean yes | Likely yes | Likely yes | Unknown | Unknown | Unknown | Unknown | Unknown | Likely yes | Unknown |
| Taste aversion behaviour | Likely yes | Likely yes | Likely yes | Likely yes | Lean yes | Likely yes | Likely yes | Likely yes | Likely yes | Lean no |
| Trace conditioning and pain | Unknown | Unknown | Lean yes | Lean yes | Likely yes | Unknown | Unknown | Unknown | Unknown | Unknown |
| Valuing behaviour | Likely yes | Lean yes | Unknown | Unknown | Lean yes | Unknown | Lean yes | Lean yes | Lean yes | Unknown |
| Brain mass | 134.5 g | 3.22 g | 0.365 g | 0.735 g | Varies | Unknown | Unknown | Unknown | 0.00302 g | Unknown |
| Brain mass to body mass ratio | 1:125 g | 1:12 g | 0.45:195 g g | 1.63 × 10−3 ± 9.12 × 10−6 | 1 g : 1 kg (Log-log plots) | Unknown | Unknown | Unknown | 1:7 | Unknown |
| Brain volume | 95077 mm3 | 730 mm3 | 0.58-3 mm3 | 0.58-3 mm3 | 12.6-926.2 mm3 | Unknown | 4.05 mm3 | Unknown | 0.60 - 1.86 mm3 | Unknown |
| Critical flicker-fusion frequency in Hz | 70 - 80 Hz | 20 - 105 Hz | 72 Hz | 72 Hz | 30 - 60 Hz | 20 - 160 Hz | 14 Hz | 50 - 60 Hz | 110 Hz | Unknown |
| Encephalization quotient | 0.58 | 0.35-2.98 | 0.37 | 0.37 | 1.23 | Unknown | Unknown | Unknown | Unknown | Unknown |
| Neuron packing density | 1.2 | 0.46. | Unknown | Unknown | Unknown | Unknown | Unknown | Unknown | Unknown | Unknown |
| Total neurons | 320-430 mio | 220 mio | 8-13 mio | 8-13 mio | 500 mio | 0.08 mio | 0.1 mio | 0.08 mio | 0.02-1.1 mio | 0.86 mio |
| Working memory load | Likely yes | Likely yes | Lean yes | Lean yes | Likely yes | Unknown | Unknown | Lean yes | Likely yes | Lean yes |

**Table S2. Example references with 90 proxies across 10 taxa. Blank spaces reflect ‘unknown’, i.e. no existing published studies found. One linked citation is provided where applicable as an example (in the full review, some taxa/ proxies identified multiple relevant references, also see Fischer, 2022).**

| **Cognitive** [**Proxies**](https://docs.google.com/document/d/1B3nyY_oh2dS9gV87ITRDkEOGM-HMtwaz/edit) | **Pig** | **Chicken** | **Carp** | **Salmon** | **Octopus** | **Shrimp** | **Crab** | **Crayfish** | **Bee** | **Silkworm** |
| --- | --- | --- | --- | --- | --- | --- | --- | --- | --- | --- |
| A not B detour test | [Jansen et al., 2009](https://link.springer.com/article/10.1007/s10071-008-0191-y) | [Regolin et al. 1995](https://www.sciencedirect.com/science/article/pii/0003347295801677) |  |  |  |  |  |  | [Mirwan et al., 2015](https://agris.fao.org/agris-search/search.do?recordID=US201500226217) |  |
| Attention bias | [Crump et al., 2018](https://www.mdpi.com/2076-2615/8/8/136/htm) | [Rodd et al. 1997](https://psycnet.apa.org/record/1997-07702-003) |  |  | [Byrne et al. 2004](https://www.sciencedirect.com/science/article/abs/pii/S000334720400301X) |  |  |  |  |  |
| Body awareness | [Broom, 2010](https://www.sciencedirect.com/science/article/pii/S0168159110001553) |  |  |  |  |  |  |  |  |  |
| Communication | [Garcia et al., 2016](https://journals.biologists.com/jeb/article/219/12/1913/15162/Honest-signaling-in-domestic-piglets-Sus-scrofa) | [Collias 1987](https://sora.unm.edu/sites/default/files/journals/condor/v089n03/p0510-p0524.pdf) | [Wilson et al. 2021](https://doi.org/10.1007/s10530-021-02502-x) | [Keenleyside & Yamamoto, 1962](https://www.jstor.org/stable/4533008) | [Hanlon & Messenger 2018](https://books.google.co.uk/books?hl=en&lr=&id=oppPDwAAQBAJ&oi=fnd&pg=PR11&dq=Hanlon+%26+Messenger+2018+communication+octopus&ots=C009P4cXME&sig=JpUcuUJQlaAtgAlhpFDi_1vIZQ8#v=onepage&q=Hanlon%20%26%20Messenger%202018%20communication%20octopus&f=false) | [Bose et al. 2017](https://www.ncbi.nlm.nih.gov/pmc/articles/PMC5490968/) | [Gleeson 1991](https://www.degruyter.com/document/doi/10.7312/baue90796-003/html) | [Kubec et al. 2019](https://www.sciencedirect.com/science/article/pii/S0044523118301141) | [Kerr, 1994](https://psycnet.apa.org/record/1994-24442-001) | [Sandler et al. 2000](https://www.cell.com/cell-chemical-biology/fulltext/S1074-5521(00)00078-8?_returnURL=https%3A%2F%2Flinkinghub.elsevier.com%2Fretrieve%2Fpii%2FS1074552100000788%3Fshowall%3Dtrue) |
| Cooperative behaviour | [Fraser et al. 1995](https://www.sciencedirect.com/science/article/pii/0168159195006105) | [Zeng et al. 2016](https://academic.oup.com/auk/article/133/4/747/5149214?login=false) | [Tan et al 2019](https://doi.org/10.1016/j.ecoleng.2018.12.002) |  | [Diamant & Shpigel 1985](https://link.springer.com/article/10.1007/BF00002584) | [Tóth & Duffy 2005](https://www.ncbi.nlm.nih.gov/pmc/articles/PMC1629045/#:~:text=Our%20experiments%20suggest%20that%20coordinated,unable%20to%20chase%20intruders%20away.) | [Booksmythe et al. 2011](https://academic.oup.com/beheco/article/23/2/285/245065) | [Haubrock et al. 2019](https://link.springer.com/article/10.1007/s11273-019-09674-3) | [Breed et al., 2004](https://pubmed.ncbi.nlm.nih.gov/14651465/) | [Zalucki et al. 2002](https://www.annualreviews.org/doi/abs/10.1146/annurev.ento.47.091201.145220) |
| Cross-modal learning |  | [Loconsole et al. 2021](https://pubmed.ncbi.nlm.nih.gov/34280814/) | [Wang & Chittka 2011](https://doi.org/10.1068%2Fic847) |  | [Young 1991](https://www.journals.uchicago.edu/doi/abs/10.2307/1542389) |  |  |  | [Zhang et al., 2014](http://bee.jxau.edu.cn/_upload/article/95/4b/e596f019453dafb98fcc1967b4f1/9624d8b1-4f3c-431b-bd7b-88f33cfb18b7.pdf) | [Wessnitze & Webb 2006](https://iopscience.iop.org/article/10.1088/1748-3182/1/3/001/meta) |
| Experience projection | [Held et al., 2000](https://www.sciencedirect.com/science/article/pii/S0003347299913222) |  |  |  |  |  |  |  |  |  |
| Gratification delay | Sommer et al., 2016 | [Abeyesinghe et al. 2005](https://www.researchgate.net/publication/236168065_Can_domestic_fowl_Gallus_gallus_domesticus_show_self-control) |  |  |  |  |  |  |  |  |
| Individual differences / "personality" | [O’Malley et al. 2019](https://pdf.sciencedirectassets.com/271155/1-s2.0-S0168159119X00106/1-s2.0-S0168159119300826/am.pdf?X-Amz-Security-Token=IQoJb3JpZ2luX2VjEAsaCXVzLWVhc3QtMSJGMEQCIC39NsXdo7o%2F55rnHOvHt4ZScXx8dFn8MGfv2c4rt3uwAiBT%2BCPkcOYNLP96LfzYWXXuYwKkWTO1YQ5Gy33W%2BbrK8irVBAj0%2F%2F%2F%2F%2F%2F%2F%2F%2F%2F8BEAUaDDA1OTAwMzU0Njg2NSIMfsDtattVmkHUrqnaKqkEBmHh8rQyATqG9PlbFIHJMDqIUAKDVOGvo6M%2BlQ%2BE71oQ%2Ft0D70z0LUo0p4bzHi933RFF65338pm1%2FpS41dgKOQD9IVX9WLpZzzfrQvk0HDBYQxQRtOyiCUP3Jky1gl2uoADZPKiVkaqoVAq33LHY7nxNtC4vBw0gEd%2BFP3EEomH6ndU5eg5s9OM04eUahJT1SExpEO6ZZnzdRUR8PbBlNa6uZUM%2BArekXylpYWM7JFUB1PFjcCGVABQ3ZMX3oWVqFG7txDBQxfpMzgU1G4sgGdBB9mlrELJ2x%2BaGT2NZ4TWpKjrqKF5mfasO76CeOtD0vo2Jz9vdLmzThgR2bJYwj%2BrIHMJ8iJt5jL5GwzQq7aLgmagMcFNgbN3wTWSRJZF6faz9AyJJyzumOHD097R2C6iXNJnRFO71z4sPgyhQ7H2Y%2BMySJwaOstHGKFc8xpHFnNw18GGr0aKMPDZeIyp3jypmIwX5%2FUVEfodM81Bb5lFV7gtMNlUW%2BxBJFkbZUXnNwgL5ZxIWawVBYv6KSCCJXR25WGXmEj%2B7xzqf68ZTVtkTGgifDKwQVvDY5VnouJD4nsA%2Bv%2F9slsFcynL1l%2FfUqgrmIyt4%2BjHIPGfZns5WpToCumyA8AhIgjCsGbPtOUoexnXvQGvvuGFwxqaXS6BjO%2BLXln5fi2n1f7BCnR4oGXL3Of%2BkPeropQLzCrT19bGESVXc%2BeChs4eDuPtR6BIskSQi6%2FuZWVkC%2FzDG45qbBjqqATChJCZOGIoZ6v04mr%2BBZL0veNw0Gl%2BdFLJjE2oP8MTIXsCJDcOcuPgxtN92Q4QLZNuw6kbsXuROWzuV9Oek5cMgI09yh6SoMnUxgmegIZxuauXuUMNiS50AUEF%2BEI8X3NbZ%2BrCnWGs0wudjJAJ33ZWfd%2F9nm9WkzDMy%2Bpe%2FdaIyueKnGSp6bKXaq8FlW87tEKxia3ywlHvdyzGhx9sgHZBYJFCotiLZh%2B1%2F&X-Amz-Algorithm=AWS4-HMAC-SHA256&X-Amz-Date=20221105T201013Z&X-Amz-SignedHeaders=host&X-Amz-Expires=300&X-Amz-Credential=ASIAQ3PHCVTYZIEDSDLH%2F20221105%2Fus-east-1%2Fs3%2Faws4_request&X-Amz-Signature=dd06d7d499b93eed0e34f6302b26fca19f3529a08a9e65754d54c2f91690ac0d&hash=96c7d2b0112e322bc7ecc68b811424b9278c4172cdf858cb0bd26d91c8026d9a&host=68042c943591013ac2b2430a89b270f6af2c76d8dfd086a07176afe7c76c2c61&pii=S0168159119300826&tid=pdf-2f200fde-e2f1-4e93-a223-b1b9e3a8d034&sid=034b886a5c6ef047437a8c39ecd62fde9b7egxrqb&type=client) | [Favati et al. 2014](https://journals.plos.org/plosone/article?id=10.1371/journal.pone.0103535) | [Huntingford et al. 2010](https://onlinelibrary.wiley.com/doi/10.1111/j.1095-8649.2010.02582.x) | [Vaz-Serrano et al., 2011](https://www.sciencedirect.com/science/article/abs/pii/S0031938411000898?via%3Dihub) | [Mather & Anderson 1993](https://psycnet.apa.org/record/1994-00613-001) | [Bardara et al. 2021](https://www.sciencedirect.com/science/article/pii/S0044848620323760?via%3Dihub#f0005) | [Su et al. 2019](https://journals.biologists.com/jeb/article/222/3/jeb188706/20741/Agonistic-behaviour-and-energy-metabolism-of-bold) | [Zhao & Feng 2015](https://academic.oup.com/cz/article/61/6/966/1800555?login=false) | [Muller et al., 2010](https://www.sciencedirect.com/science/article/pii/S0003347210003854) | [Obara & Tamazawa 1982](https://resjournals.onlinelibrary.wiley.com/doi/10.1111/j.1365-3032.1982.tb00320.x) |
| Inhibitory control | [Zebunke et al., 2018](https://www.frontiersin.org/articles/10.3389/fpsyg.2018.02099/full) | [Wascher et al. 2021](https://royalsocietypublishing.org/doi/abs/10.1098/rsos.210504) | [Lucon-Xiccato & Bertolucci 2020](https://doi.org/10.1111/jfb.14380) |  |  |  |  |  | [Mayack and Naug, 2015](https://royalsocietypublishing.org/doi/full/10.1098/rsbl.2014.0820) |  |
| Judgment bias | [Düpjan et al., 2013](https://www.sciencedirect.com/science/article/pii/S1558787813001329) | [Crump et al. 2018](https://pubmed.ncbi.nlm.nih.gov/30087230/) | [Espigares et al. 2021](https://royalsocietypublishing.org/doi/full/10.1098/rsbl.2020.0745) |  | [Schnell & Vallortigara 2019](https://www.wellbeingintlstudiesrepository.org/cgi/viewcontent.cgi?article=1502&context=animsent) |  |  | [Bacqué-Cazenave et al., 2017](https://www.nature.com/articles/srep39935?origin=ppub) | [Bateson et al., 2011](https://pubmed.ncbi.nlm.nih.gov/21636277/) |  |
| Memory bias |  |  |  |  |  |  |  |  |  |  |
| Mental Time Travel | [Kouwenberg et al. 2009](https://www.sciencedirect.com/science/article/abs/pii/S0168159109000240) | [Marino 2017](https://link.springer.com/article/10.1007/s10071-016-1064-4/) | [Hamilton et al., 2016](https://link.springer.com/article/10.1007/s10071-016-1014-1) |  | [Jozet-Alves et al. 2013](https://pubmed.ncbi.nlm.nih.gov/24309275/) |  |  |  |  |  |
| Mirror mark recognition | [Broom et al., 2009](https://psycnet.apa.org/record/2009-20075-003) | [Gallup Jr, 1975](https://scholar.google.cz/scholar?q=(Gallup,+G.G.+(1975)&hl=en&as_sdt=0&as_vis=1&oi=scholart) |  |  | [Amodio & Fiorito 2022](https://pubmed.ncbi.nlm.nih.gov/36111145/) |  |  |  |  |  |
| Motivational trade-off | [Kratzer, 1969](https://academic.oup.com/jas/article-abstract/28/2/175/4698313?redirectedFrom=fulltext) | [Appleby et al., 2004](https://books.google.co.uk/books?hl=en&lr=&id=hW6BCwAAQBAJ&oi=fnd&pg=PR9&dq=appleby+2004+chicken+motivational+trade+off&ots=uVH3HXOjCZ&sig=rXlwSVLfo77LdgNPA_ZQ-Pdqf50#v=onepage&q&f=false) | [Dunlop et al. 2006](https://www.sciencedirect.com/science/article/abs/pii/S0168159105001802) | [Dunlop et al. 2006](https://www.sciencedirect.com/science/article/abs/pii/S0168159105001802) | [Crook et al. 2011](https://pubmed.ncbi.nlm.nih.gov/21900465/) | [Maskrey et al. 2018](https://www.sciencedirect.com/science/article/abs/pii/S0003347218301416?via%3Dihub) | [Magee & Elwood 2013](https://journals.biologists.com/jeb/article/216/3/353/11942/Shock-avoidance-by-discrimination-learning-in-the) | [Mergler et al. 2020](https://link.springer.com/article/10.1007/s10452-020-09775-9) | Gibbons et al. 2022 | [Mir & Qamar 2018](https://link.springer.com/article/10.1007/s13744-017-0559-2#Bib1) |
| Multimodal integration | [Statham et al. 2020](https://www.nature.com/articles/s41598-020-65954-6) | [Verhaal and Luksch 2016](https://www.researchgate.net/publication/283811564_Multimodal_integration_in_behaving_chickens) | [Wang & Chittka 2011](https://doi.org/10.1068%2Fic847) |  | [Gutnick et al. 2011](https://www.sciencedirect.com/science/article/pii/S0960982211001084) | [Hebets & Rundus 2011](https://link.springer.com/chapter/10.1007/978-0-387-77101-4_17) | [Sneddon et al.2003](https://link.springer.com/article/10.1023/A:1021972412694) | [Aquiloni & Gherardi 2008](https://onlinelibrary.wiley.com/doi/10.1111/j.1365-2427.2007.01911.x) | [Ostwald et al., 2019).](https://www.sciencedirect.com/science/article/pii/S0003347219301976) | [Yamada et al 2021](https://elifesciences.org/articles/72001) |
| Navigation strategies | [Morelle et al. 201](https://onlinelibrary.wiley.com/doi/10.1111/mam.12028)5 | [Denzau et al. 2013](https://journals.biologists.com/jeb/article/216/16/3143/11569/Ontogenetic-development-of-magnetic-compass) | [Moorman, 2001](https://academic.oup.com/ilarjournal/article/42/4/292/743671) | [Dittman & Quinn, 1996](https://pubmed.ncbi.nlm.nih.gov/9317381/) | [Forsythe & Hanlon 1997](https://www.sciencedirect.com/science/article/abs/pii/S0022098196000573) | [Reaka 1980](https://www.sciencedirect.com/science/article/abs/pii/S0003347280800145) | [Keller et al. 2003](https://www.int-res.com/articles/meps2003/261/m261p217.pdf) | [Kamran & Moore 2015](https://onlinelibrary.wiley.com/doi/10.1111/eth.12392) | [Chittka & Geiger 1995](http://chittkalab.sbcs.qmul.ac.uk/1995/ChittkaGeiger_AnBehav95.pdf) | [Namiki et al. 2018](https://www.nature.com/articles/s41598-018-27954-5) |
| Numerical cognition |  | [Rugani et al. 2015](https://www.researchgate.net/publication/317719814_Number-space_mapping_in_the_newborn_chick_resembles_humans'_mental_number_line) | [Xiong et al., 2018](https://doi.org/10.1007/s10071-018-1214-y) |  |  |  |  |  | Eckert et al., 2021 |  |
| Object permanence | [Nawroth et al., 2013](https://aab.copernicus.org/articles/56/861/2013/) | [Regolin et al. 1995](https://www.sciencedirect.com/science/article/pii/S0003347285702321) | [Sovrano et al. 2018).](https://www.frontiersin.org/articles/10.3389/fpsyg.2018.02341/full) | [Sovrano et al. 2018).](https://www.frontiersin.org/articles/10.3389/fpsyg.2018.02341/full) |  |  |  |  |  |  |
| Perspective taking | [Held et al., 2005](https://link.springer.com/article/10.1007/s10071-004-0242-y) |  |  |  |  |  |  |  |  |  |
| Physical reasoning | [Albiach-Serrano et al. 2012](https://www.sciencedirect.com/science/article/pii/S0168159112002225?via%3Dihub) | [Chiandetti and Vallortigara 2011](https://www.jstor.org/stable/41314975) |  |  | [Finn et al. 2009](https://www.sciencedirect.com/science/article/pii/S0960982209019149) |  |  |  |  |  |
| Problem-solving | [Pérez Fraga et al., 2021](https://link.springer.com/article/10.1007%2Fs10071-020-01410-2) | [Daisley et al. 2010](https://pubmed.ncbi.nlm.nih.gov/20178037/) |  |  | [Fiorito et al. 1998](https://pubmed.ncbi.nlm.nih.gov/18953584/) | [Duffield et al. 2015](https://journals.plos.org/plosone/article?id=10.1371/journal.pone.0139050) |  |  | [Loukola et al. 2017](https://pubmed.ncbi.nlm.nih.gov/28232576/) |  |
| Response-slowing |  |  |  |  |  |  |  |  |  |  |
| Responses to novelty | [Lewis et al., 2008](https://www.sciencedirect.com/science/article/pii/S187114130800070X?via%3Dihub) | [Garnham and Løvlie 2018](https://www.ncbi.nlm.nih.gov/pmc/articles/PMC5791031/) | [Lucon-Xiccato and Dadda, 2014](https://doi.org/10.1016/j.beproc.2014.03.010) | [Wilson & Stevens 2005](https://onlinelibrary.wiley.com/doi/abs/10.1111/j.1439-0310.2005.01110.x) | [Mather & Anderson 1993](https://psycnet.apa.org/doiLanding?doi=10.1037%2F0735-7036.107.3.336) |  |  | [Shuranova et al. 2005](https://academic.oup.com/jcb/article/25/3/488/2670536?login=false) | [Muller et al., 2010](https://www.sciencedirect.com/science/article/pii/S0003347210003854) |  |
| Reversal learning | [Bolhuis et al., 2004](https://www.sciencedirect.com/science/article/pii/S0166432803003887?via%3Dihub) | [Wascher et al. 2021](https://royalsocietypublishing.org/doi/abs/10.1098/rsos.210504) | [Kuroda et al. 2017](https://pubmed.ncbi.nlm.nih.gov/28633953/) | [Ruiz-Gomez et al. 2011](https://pubmed.ncbi.nlm.nih.gov/21130105/) | [Boycott & Young, 1957](https://royalsocietypublishing.org/doi/10.1098/rspb.1957.0023) | [Ventura & Mattei 1977](https://reader.elsevier.com/reader/sd/pii/S0091677377906447?token=65D106D57A42A4FC0AAF0F274EED3B8AFAA9B6FC6F357347FF436E42A70FB84315E17BE5D1D20EF303121F8B73FD64E5&originRegion=eu-west-1&originCreation=20220403195753) | [Abramson & Feinman 1990](https://www.sciencedirect.com/science/article/pii/003193849090311Q) | [Tierney et al. 201](https://link.springer.com/article/10.3758/s13420-018-0345-y)9 | [Raine & Chittka 2012](https://pubmed.ncbi.nlm.nih.gov/23028779/) | [Rodrigues et al. 2010](https://onlinelibrary.wiley.com/doi/abs/10.1111/j.1439-0310.2009.01737.x) |
| Reverse reward test |  |  |  |  | [Bublitz et al., 2017](https://www.frontiersin.org/articles/10.3389/fphys.2017.00054/full) |  |  |  |  |  |
| Shared intentionality |  |  |  |  |  |  |  |  |  |  |
| Social learning | [Bolhuis et al., 2011](https://www.sciencedirect.com/science/article/pii/S0003347211002417) | [Nicol and Pope 1994](http://psycnet.apa.org/record/1995-00506-001) | [Bajer et al. 2010](https://d1wqtxts1xzle7.cloudfront.net/44233967/Cognitive_aspects_of_food_searching_beha20160330-21357-iu4axl-with-cover-page-v2.pdf?Expires=1667725282&Signature=WFKGwc6b0HEeTuhVpFW8-VTzGPXhMjE1pJinPRKVZWeCLoK2Sy-3VJzXmKLyQCrCh70kVIEdbsdRA-OORQyTSgo3-VrRncCY8rzolLm9rnAlpTxzGChD46szfbH6fO-QZ-x72g4qBmK-OLyUquxkEoZE8u9URs9jkZ2flnlGt24Fq7nOx51TN1UBFb4Gvp3GE-DxSwKUMwidMzKWqu2z2P6m5Bd~6id62LVUzfKhof~I1Hie1P~jHh4mJUHbpDdxGWJU3E8S6jZUBErOyRIFgUbtKKuI9kS00JqhsWOIIKVB1L1n3-tkeyOZZi~eEzRP5O9N6zsAB6S44HfprWY6zQ__&Key-Pair-Id=APKAJLOHF5GGSLRBV4ZA) | [Brown & Laland 2002](https://onlinelibrary.wiley.com/doi/abs/10.1111/j.1095-8649.2002.tb01857.x) | [Fiorito & Scotto 1992](https://psycnet.apa.org/record/1992-37971-001) |  |  | [Jiménez-Morales et al. 2018](https://www.sciencedirect.com/science/article/pii/S1074742717302022) | [Alem et al., 2016](https://journals.plos.org/plosbiology/article?id=10.1371/journal.pbio.1002589) |  |
| Socio-spatial cognition | [Arts et al. 2009](https://www.sciencedirect.com/science/article/pii/S0166432809003696) | [Campbell et al. 2018](https://www.mdpi.com/2076-2615/8/2/26) | [Ghosal et al., 2016](https://journals.plos.org/plosone/article?id=10.1371/journal.pone.0157174) | [Dittman & Quinn, 1996](https://doi.org/10.1242/jeb.199.1.83) |  |  |  |  | [Mirwan & Kevan, 2015](https://psycnet.apa.org/record/2015-22252-001) |  |
| Symbolic representation of the world |  | [Freire 2020](https://researchoutput.csu.edu.au/en/publications/understanding-chicken-learning-and-cognition-and-implications-for) |  |  |  |  |  |  |  |  |
| Temporal/ spatial discounting | [Ferguson et al., 2009](https://pubmed.ncbi.nlm.nih.gov/18804519/) | [Abeyesinghe et al. 2005](https://www.researchgate.net/publication/236168065_Can_domestic_fowl_Gallus_gallus_domesticus_show_self-control) |  |  |  |  |  |  | [Cheng et al., 2002](https://link.springer.com/article/10.3758/BF03196280) |  |
| Theory of mind | [Held et al., 2005](https://link.springer.com/article/10.1007/s10071-004-0242-y) |  |  |  |  |  |  |  |  |  |
| Tool use | [Root-Bernstein et al., 2019](https://www.sciencedirect.com/science/article/pii/S1616504719300333) |  |  |  | [Finn et al. 2009](https://www.sciencedirect.com/science/article/pii/S0960982209019149) |  | [Thanh et al, 2005](https://link.springer.com/article/10.1007/s00227-005-0017-2) |  | [Loukola et al., 2017](https://www.science.org/doi/10.1126/science.aag2360) |  |
| Transitive inference | [Vasconcelos, 2008](https://www.sciencedirect.com/science/article/pii/S0376635708000818?via%3Dihub) | [Hogue et al. 1996](http://cogprints.org/1939/3/hogue-beaugrand-lague-1996.pdf) |  | [White & Gowan 2013](https://doi.org/10.1093/beheco/ars136) |  |  |  |  | [Benard & Giurfa, 2004](https://psycnet.apa.org/record/2004-14996-013) |  |
| Uncertainty monitoring |  |  |  |  |  | [Bool et al, 2011](https://www.wellbeingintlstudiesrepository.org/cgi/viewcontent.cgi?article=1071&context=acwp_asie) |  | [Tierney and Andrews, 2013](https://link.springer.com/article/10.1007/s10071-012-0547-1) | [Perry & Barron, 2013](https://www.discovermagazine.com/planet-earth/honey-bees-selectively-avoid-difficult-choices) |  |
| Anxiety-like behaviour | [Norscia et al., 2021](https://pubmed.ncbi.nlm.nih.gov/33665219/) | [Rodd et al., 1997](https://www.sciencedirect.com/science/article/pii/S0023969097909528/pdf?md5=39cf9fcafea3c40d2957424e15ad86f2&pid=1-s2.0-S0023969097909528-main.pdf) | [Todorova et al. 2015](http://www.uni-sz.bg/tsj/Vol.%2013,%202015,%20Suppl.%202,%20Series%20Biomedical%20Sciences/AF/AF/Ecology/K.%20Todorova.pdf) | [Geller & Brady 1961](https://www.science.org/doi/10.1126/science.133.3458.1080) | [Bennett & Toll 2011](https://www.ncbi.nlm.nih.gov/pmc/articles/PMC3228935/#:~:text=Intramantle%20inking%20and%20distributed%20rigor,animal%20to%20a%20healthy%20state.) | [Takahashi 2022](https://link.springer.com/article/10.1007/s10071-021-01555-8#citeas) | [Wilson et al 2021](https://www.tandfonline.com/doi/abs/10.1080/10236244.2021.1923369) | [Fossat et al 2014](https://www.science.org/doi/10.1126/science.1248811) | [Bateson et al., 2011](https://www.sciencedirect.com/science/article/pii/S0960982211005446#bib31) |  |
| Associative learning from pain |  | [Danbury et al. 2000](https://bvajournals.onlinelibrary.wiley.com/doi/pdfdirect/10.1136/vr.146.11.307) | [Yoshida & Hirano 2010](https://behavioralandbrainfunctions.biomedcentral.com/articles/10.1186/1744-9081-6-20) | [Dunlop et al. 2006](https://doi.org/10.1016/j.applanim.2005.06.018) | [Crook 2021](https://www.cell.com/iscience/pdf/S2589-0042(21)00197-8.pdf) |  | [Magee & Elwood 2013](https://journals.biologists.com/jeb/article/216/3/353/11942/Shock-avoidance-by-discrimination-learning-in-the) |  | [Kirkerud et al., 2017](https://www.frontiersin.org/articles/10.3389/fnbeh.2017.00094/full) |  |
| Boredom-like behaviour | [Wemelsfelder, 1985](https://link.springer.com/chapter/10.1007/978-94-009-4998-0_8) | [Renner and Rosenzweig, 1987](https://link.springer.com/book/10.1007/978-1-4612-4766-1) |  |  | [Anderson & Mather 2007](https://psycnet.apa.org/buy/2007-11961-008) |  |  |  |  |  |
| Concept of death |  |  |  |  |  |  |  |  |  |  |
| Curiosity-like behaviour |  |  | [Murphy & Pitcher, 1991](https://doi.org/10.1111/j.1439-0310.1991.tb00285.x) |  | [Mather 1994](https://zslpublications.onlinelibrary.wiley.com/doi/abs/10.1111/j.1469-7998.1994.tb05270.x) |  |  |  |  |  |
| Depression-like behaviour | [Sadler et al., 2014](https://www.ingentaconnect.com/content/ufaw/aw/2014/00000023/00000002/art00003;jsessionid=27346d4ythdoh.x-ic-live-02) | [Sufka et al., 2006](https://pubmed.ncbi.nlm.nih.gov/17110794/) | [Orjes 2015](https://lup.lub.lu.se/luur/download?func=downloadFile&recordOId=7862046&fileOId=7862056) | [Shapouri 2020](https://nmbu.brage.unit.no/nmbu-xmlui/handle/11250/2725049) |  |  |  |  |  |  |
| Disgust-like behaviour | [Jones et al., 2000](https://www.sciencedirect.com/science/article/pii/S0168159100001490) | [Johnston et al. 1998](https://www.semanticscholar.org/paper/Observation-learning-in-day-old-chicks-using-a-Johnston-Burne/af409fa36fafd03374396a1568c94e6e972368af) |  |  |  |  |  |  |  |  |
| Displacement behaviour | [Marcet-Rius et al., 2019](https://www.sciencedirect.com/science/article/pii/S1871141319301350) | [Ferreira et al. 2021](https://www.tandfonline.com/doi/full/10.1080/00439339.2021.1924920) |  |  |  | [Bardera et al. 2019](https://www.sciencedirect.com/science/article/abs/pii/S0044848619312220) |  |  | [Rao et al., 2019](https://experts.umn.edu/en/publications/remarkable-long-distance-returns-to-a-forage-patch-by-artificiall) |  |
| Effects on exploratory behaviour from pain | [Kluivers-Poodt et al., 2013](https://www.sciencedirect.com/science/article/pii/S1751731113000086) | [Gentle et al. 1990](https://www.sciencedirect.com/science/article/pii/0168159190900145) | [Deakin et al. 2019](https://doi.org/10.3390/fishes4010008) | [Sneddon et al. 2003](https://onlinelibrary.wiley.com/doi/abs/10.1046/j.1095-8649.2003.00084.x) | [Ross 1971](https://www.nature.com/articles/230401a0) |  |  |  |  |  |
| Embarrassment-like behaviour |  |  |  |  |  |  |  |  |  |  |
| Emotional contagion | [Reimert et al., 2013](https://www.sciencedirect.com/science/article/pii/S0031938412003447) | [Edgar & Nicol 2018](https://www.nature.com/articles/s41598-018-28923-8) | [Oliveira et al. 2017](https://doi.org/10.7717/peerj.3739) |  |  |  |  |  |  |  |
| Emotional reactions to learning | [Reimert et al., 2013](https://www.sciencedirect.com/science/article/pii/S0031938412003447) | [Khune et al. 2013](https://www.sciencedirect.com/science/article/pii/S0168159113002268?via%3Dihub) |  |  | [Ross 1971](https://www.nature.com/articles/230401a0) |  |  |  |  |  |
| Exhaustion-like behaviour | [Bradshaw et al., 1996](https://www.cambridge.org/core/journals/animal-science/article/abs/behavioural-and-hormonal-responses-of-pigs-during-transport-effect-of-mixing-and-duration-of-journey/02D1A830ED8B3961BEF7118B1D3A9155) | [Dayyani and Bakhtiari, 2013](http://www.ijabbr.com/article_7968.html) | [Knudsen & Jensen, 1998](https://doi.org/10.1016/S1095-6433(97)00435-2) | [Milligan, 1996](https://doi.org/10.1016/0300-9629(95)02060-8) | [Franklin et al. 2017](https://onlinelibrary.wiley.com/doi/full/10.1111/jeb.13063) | [Robles-Romo et al. 2016](https://www.sciencedirect.com/science/article/abs/pii/S0022098116300156?via%3Dihub) |  | [Misra et al. 1996](https://www.astm.org/stp11719s.html) |  | [Bura et al. 2011](https://journals.biologists.com/jeb/article/214/1/30/10205/Whistling-in-caterpillars-Amorpha-juglandis) |
| Fear-like behaviour | [Arroyo et al., 2016).](https://www.sciencedirect.com/science/article/pii/S0031938416303900) | [Duncan and Petherick 1991](https://pubmed.ncbi.nlm.nih.gov/1808195/) | [Stabell et al. 2010](https://doi.org/10.1007/s10641-010-9707-9) | [Geller & Brady 1961](https://www.science.org/doi/10.1126/science.133.3458.1080) | [Bennett & Toll 2011](https://www.ncbi.nlm.nih.gov/pmc/articles/PMC3228935/#:~:text=Intramantle%20inking%20and%20distributed%20rigor,animal%20to%20a%20healthy%20state.) | [Takahashi 2022](https://link.springer.com/article/10.1007/s10071-021-01555-8#citeas) | [Wilson et al 2021](https://www.tandfonline.com/doi/abs/10.1080/10236244.2021.1923369) | [Wood & Moore 2020](https://cdnsciencepub.com/doi/abs/10.1139/cjz-2019-0089) | [Tan et al. 2013](https://journals.plos.org/plosone/article?id=10.1371/journal.pone.0075841) |  |
| Flexible self-protective behaviour | [Bracke, 2011](https://www.sciencedirect.com/science/article/pii/S0168159111000219) | [Duncan et al. 1989](https://www.tandfonline.com/doi/pdf/10.1080/00071668908417172) | Reilly et al. 2008 | [Reilly et al. 2008](https://doi.org/10.1016/j.applanim.2008.01.016) | [Alupay et al. 2014](https://pubmed.ncbi.nlm.nih.gov/24239646/) | [Bauer 1981](https://academic.oup.com/jcb/article-abstract/1/2/153/2327452?redirectedFrom=fulltext#no-access-message) | [McCambridge et al. 2016](https://bioone.org/journals/Journal-of-Shellfish-Research/volume-35/issue-4/035.035.0426/Effects-of-Autotomy-Compared-to-Manual-Declawing-on-Contests-between/10.2983/035.035.0426.short) | [Puri & Faulkes 2010](https://journals.plos.org/plosone/article?id=10.1371/journal.pone.0010244) | [Breed et al., 1990](https://www.jstor.org/stable/4600497) | [Walters et al. 2001](https://pubmed.ncbi.nlm.nih.gov/11171298/) |
| Friendship-like behaviour | [Ewbank and Meese, 1971](https://www.cambridge.org/core/journals/animal-science/article/abs/aggressive-behaviour-in-groups-of-domesticated-pigs-on-removal-and-return-of-individuals/5571226ED68E7B1F863D26B1F15192EB) |  | [Miller & Gerlai, 2012](https://doi.org/10.1371/journal.pone.0048865) |  |  |  |  |  |  |  |
| Helping behaviour | [Hemsworth et al., 2011](https://www.sciencedirect.com/science/article/pii/S0376635711000751) | [Nicol and Pope 1994](https://psycnet.apa.org/record/1995-00506-001) |  |  |  |  |  | [Mathews 2011](https://brill.com/view/journals/beh/148/1/article-p71_4.xml) | [Rueppell et al., 2010](https://onlinelibrary.wiley.com/doi/10.1111/j.1420-9101.2010.02022.x) |  |
| Hyperalgesia | [Steagall et al., 2021](https://www.mdpi.com/2076-2615/11/6/1483) | [Sufka and Hughes 1990](https://psycnet.apa.org/record/1990-19187-001) |  |  | [Aluplay et al. 2014](https://pubmed.ncbi.nlm.nih.gov/24239646/) |  |  |  |  | [Walters et al. 2001](https://pubmed.ncbi.nlm.nih.gov/11171298/) |
| Jealousy-like behaviour |  |  |  |  |  |  |  |  |  |  |
| Joy-like behaviour |  |  | [Graham et al. 2018](https://doi.org/10.1016/j.applanim.2018.02.005) |  | [Mather & Anderson 1999](https://psycnet.apa.org/record/1999-03911-013) |  |  |  |  |  |
| Liking-wanting dissociation |  |  |  |  |  |  |  |  |  |  |
| Loneliness-like behaviour | [van der Staay et al., 2016](https://www.ncbi.nlm.nih.gov/pmc/articles/PMC4707236/) | [Lehr, 1989](https://pubmed.ncbi.nlm.nih.gov/2498920/) | [Kittilsen 2013](https://doi.org/10.1016/j.beproc.2013.09.002) | [Kittilsen 2013](https://doi.org/10.1016/j.beproc.2013.09.002) |  |  |  |  |  |  |
| Love-like behaviour |  |  |  |  |  |  |  |  |  |  |
| Maternal response to offspring distress | [Hutson et al., 1992](https://www.sciencedirect.com/science/article/pii/S0168159105800917) | [Edgar et al. 2011](https://royalsocietypublishing.org/doi/10.1098/rspb.2010.2701) |  |  |  | [da Costa & Fransozo 2004](https://link.springer.com/article/10.1007/s10750-004-6410-x) |  |  |  | [Banno et al. 2010](https://www.researchgate.net/publication/44613626_The_Silkworm-An_Attractive_BioResource_Supplied_by_Japan) |
| Mourning-like behaviour |  |  |  |  |  |  |  |  |  |  |
| Panic-like behaviour | [Temple et al., 2011](https://www.sciencedirect.com/science/article/pii/S0168159111000323) | [Gallup and Suarez, 1980](https://www.sciencedirect.com/science/article/abs/pii/S0003347280800455) | [Vøllestad et al. 2004](https://onlinelibrary.wiley.com/doi/10.1111/j.1600-0633.2004.00048.x+) | [Gismervik et al., 2019](https://www.semanticscholar.org/paper/Thermal-injuries-in-Atlantic-salmon-in-a-pilot-Gismervik-G%C3%A5snes/40785d425e66a245934624f135d6f9af4c07db96) | [Hanlon & Messenger 2018](https://archive.org/details/Cephalopoda-behaviour-2018) |  |  |  |  |  |
| Parental care | [Drake et al., 2008](https://link.springer.com/article/10.1007/s00265-007-0418-y) | [Edgar et al. 2016](https://www.mdpi.com/2076-2615/6/1/2) | [Uusi-Heikkilä et al. 2012](https://link.springer.com/article/10.1007/s10641-011-9937-5) | [Baxter & MacPhail 1999](https://cdnsciencepub.com/doi/10.1139/z99-090) |  | [da Costa & Fransozo 2004](https://link.springer.com/article/10.1007/s10750-004-6410-x) | [Ruiz-Tagle et al. 2002](https://www.sciencedirect.com/science/article/pii/S0022098102002381?via%3Dihub#BIB8) | [Mathews 2011](https://brill.com/view/journals/beh/148/1/article-p71_4.xml) | [Hogendoorn et al., 2001](https://www.semanticscholar.org/paper/Extended-alloparental-care-in-the-almost-solitary-Hogendoorn-Watiniasih/7ea6128a11f11bb96e29ed7901700b41e399fa64) | [Banno et al. 2010](https://www.researchgate.net/publication/44613626_The_Silkworm-An_Attractive_BioResource_Supplied_by_Japan) |
| Play behaviour | Donaldson et al., 2002 | [Dawson and Siegel 1967](https://vtechworks.lib.vt.edu/handle/10919/42826) | [Burghardt, 2006](https://doi.org/10.7551/mitpress/3229.003.0018) | [Fagen, 2017](https://doi.org/10.3390/ani7060042) | [Kuba et al. 2006](https://pubmed.ncbi.nlm.nih.gov/16893255/) |  |  |  | [Loukola et al. 2017](https://pubmed.ncbi.nlm.nih.gov/28232576/) |  |
| Play vocalization | [Friel et al., 2019](https://www.nature.com/articles/s41598-019-38514-w) |  |  |  |  |  |  |  |  |  |
| Prioritizes pain response in relevant context | [Rushen et al., 1993](https://www.sciencedirect.com/science/article/abs/pii/003193849390203R) | [Wylie and Gentle 1998](https://doi.org/10.1016/S0031-9384(98)00020-1) | [Deakin et al. 2019](https://awionline.org/lab-animal-search/deakin-g-buckley-j-alzubi-h-s-et-al-2019-automated-monitoring-behaviour-zebrafish) | [Sneddon et al. 2003](https://doi.org/10.1067/S1526-5900(03)00717-X) | [Crook 2021](https://www.cell.com/iscience/pdf/S2589-0042(21)00197-8.pdf) | [Taylor et al. 2004](https://www.sciencedirect.com/science/article/abs/pii/S0044848603006616) |  |  |  |  |
| PTSD-like behaviour | [Warriss](https://www.ingentaconnect.com/content/ufaw/aw/1998/00000007/00000004/art00002) et al. 1998 |  | [Yang et al. 2020](https://doi.org/10.1016/j.bbr.2020.112644) |  |  |  |  |  |  |  |
| Relief learning | [Imfeld-Mueller et al., 2011](https://www.sciencedirect.com/science/article/pii/S016815911100058X) |  | [Sneddon et al. 2014](https://doi.org/10.1016/j.anbehav.2014.09.007) |  | [Crook 2020](https://www.biorxiv.org/content/10.1101/2020.08.23.263426v1#:~:text=In%20conditioned%20place%20preference%20assays,invertebrate%20shown%20to%20experience%20pain.) | [Bool et al. 2011](https://www.publish.csiro.au/mf/MF11140) | [Magee and Elwood, 2013](https://journals.biologists.com/jeb/article/216/3/353/11942/Shock-avoidance-by-discrimination-learning-in-the) | [Okada et al., 2021](https://journals.biologists.com/jeb/article/224/6/jeb242180/237927/Aversive-operant-conditioning-alters-the) | [Yarali et al., 2012](https://journals.plos.org/plosone/article?id=10.1371/journal.pone.0032885) |  |
| Rescue behaviour | [Masilkova et al., 2021](https://www.nature.com/articles/s41598-021-95682-4#Tab1) | [Hammers and Brouwer 2017](https://brill.com/view/journals/beh/154/4/article-p403_2.xml) |  |  |  |  |  |  |  |  |
| Response modified by painkillers | [McGlone and Hellman, 1988](https://www.depts.ttu.edu/animalwelfare/research/pigcastration/documents/CastrationandbehaviorMcGloneandHellman1988.pdf) | [Singh et al. 2017](https://www.sciencedirect.com/science/article/pii/S1467298717300296) | [Chervova & Lapshin 2000](https://www.researchgate.net/profile/Dmitry-Lapshin-2/publication/12118861_Opioid_Modulation_of_Pain_Threshold_in_Fish/links/5a98008145851535bcdf955e/Opioid-Modulation-of-Pain-Threshold-in-Fish.pdf) | [Nordgreen et al. 2013](https://doi.org/10.1016/j.applanim.2013.03.002) | [Butler-Struben et al. 2018](https://pubmed.ncbi.nlm.nih.gov/29515454/) | [Taylor et al. 2004](https://www.sciencedirect.com/science/article/pii/S0044848603006616) | [Barr & Elwood 2011](https://pubmed.ncbi.nlm.nih.gov/21324350/) | [Buřič et al. 2018](https://www.sciencedirect.com/science/article/pii/S0166445X18304259?via%3Dihub) | [Groening et al., 2017](https://www.nature.com/articles/srep45825) |  |
| Reward based learning | [Hemsworth et al., 1996](https://www.sciencedirect.com/science/article/pii/0168159196010659) | [Ferreira et al. 2020](https://www.researchgate.net/publication/342963693_Range_use_is_related_to_free-range_broiler_chickens'_behavioral_responses_during_food_and_social_conditioned_place_preference_tests) | [Wright & Eastcott 1982](https://doi.org/10.1111/j.1095-8649.1982.tb03972.x) | [Paspatis & Boujard 1996](https://www.sciencedirect.com/science/article/abs/pii/S0044848696013361) | [Mackintosh & Mackintosh 1963](https://psycnet.apa.org/record/1965-04403-001) | [Ventura and Mattel, 1977](https://www.sciencedirect.com/science/article/abs/pii/S0091677377906447) | Davies et al 2019 | [Imeh-Nathaniel et al. 2016](https://www.sciencedirect.com/science/article/abs/pii/S0031938415301475?via%3Dihub) | [Amaya-Márquez et al., 2019](https://www.mdpi.com/2075-4450/10/11/412) | [Takahashi et al. 2021](https://zenodo.org/record/5609468#.YiILjN_P2Uk) |
| Sadness-like behaviour |  |  |  |  |  |  |  |  |  |  |
| Self-medication | [Huffman, 2003](https://www.cambridge.org/core/journals/proceedings-of-the-nutrition-society/article/animal-selfmedication-and-ethnomedicine-exploration-and-exploitation-of-the-medicinal-properties-of-plants/7F3DFBB1BEE144895048E04F0613E038) | [Danbury et al. 2000](https://pubmed.ncbi.nlm.nih.gov/10766114/) | [Darland & Dowling 2001](https://doi.org/10.1073/pnas.191380698) | [Darland & Dowling 2001](https://doi.org/10.1073/pnas.191380698) | [Crook 2020](https://www.biorxiv.org/content/10.1101/2020.08.23.263426v1#:~:text=In%20conditioned%20place%20preference%20assays,invertebrate%20shown%20to%20experience%20pain.) |  |  | [MacKay & Moore 2021](https://academic.oup.com/jcb/article/41/4/ruab057/6398516?login=true) | [Behmer, 2009](https://www.annualreviews.org/doi/abs/10.1146/annurev.ento.54.110807.090537) | [Singer et al. 2004](https://pubmed.ncbi.nlm.nih.gov/15478095/) |
| Sensory-Affective Dissociation |  | [Moore and Capretta, 1968](https://link.springer.com/article/10.3758/BF03331266) |  |  | [Crook 2020](https://www.biorxiv.org/content/10.1101/2020.08.23.263426v1#:~:text=In%20conditioned%20place%20preference%20assays,invertebrate%20shown%20to%20experience%20pain.) |  |  |  |  |  |
| Social buffering | [Puppe, 1998](https://www.sciencedirect.com/science/article/pii/S0168159198001075?casa_token=xkZLhB1rwgsAAAAA:qalbDnBCwg7oNmowFFaRizXV1IdLQaA734g3vhSInYJCyaV9CFptkb6gZvtYdYdmDixoMKDsNA) | [Edgar et al. 2016](https://www.mdpi.com/2076-2615/6/1/2) | [Faustino et al. 2017](https://doi.org/10.1038/srep44329) |  |  |  |  |  | [Poot-Báez et al., 2020](https://hal.archives-ouvertes.fr/hal-03084270/document) |  |
| Taste aversion behaviour | [Houpt et al., 1979)](https://link.springer.com/article/10.1007/BF02268964) | [Johnston et al. 1998](https://pubmed.ncbi.nlm.nih.gov/9933530/) | [Kasumyan & Morsi 1995](https://www.bouillettes-dependance-baits.com/res/site19627/res552278_taste-common-carp.pdf) | [Hughes, 1991](https://www.tandfonline.com/doi/abs/10.1577/1548-8640(1991)053%3C0015%3ACROFFS%3E2.3.CO%3B2) | [van Giesen et al. 2020](https://www.sciencedirect.com/science/article/pii/S0092867420311491) | [Bardera et al. 2020](https://www.sciencedirect.com/science/article/pii/S004484862031629X?via%3Dihub#bb0240) | [Grober 1988](https://www.sciencedirect.com/science/article/abs/pii/S0003347288800204) | [Arzuffi et al., 2000](https://www.sciencedirect.com/science/article/abs/pii/S0031938499002280) | [de Brito Sanchez, 2011](https://academic.oup.com/chemse/article/36/8/675/265934) | [Banno et al. 2010](https://www.researchgate.net/publication/44613626_The_Silkworm-An_Attractive_BioResource_Supplied_by_Japan) |
| Trace conditioning and pain |  |  | [Overmeier & Savage 1974](https://doi.org/10.1016/0014-4886(74)90031-4) | [Nordgreen et al. 2010](https://link.springer.com/article/10.1007/s10071-009-0267-3) |  |  |  |  |  |  |
| Valuing behaviour | [Held et al. 2005](https://link.springer.com/article/10.1007/s10071-004-0242-y) | [Vas et al. 2020](https://www.sciencedirect.com/science/article/pii/S0168159120302173?via%3Dihub) |  |  |  |  | Sneddon et al. 1996 | [Buric et al. 2016](https://springerplus.springeropen.com/articles/10.1186/s40064-016-3343-6) | [Sanderson et al. 2013](https://doi.apa.org/doiLanding?doi=10.1037%2Fa0032613) |  |
| Brain mass | [Minervini et al., 2016](https://journals.plos.org/plosone/article?id=10.1371/journal.pone.0157378) | [Henriksen et al. 2016](https://www.nature.com/articles/srep34031) | [Khan et al. 2022](https://doi.org/10.1590/1519-6984.242897) | [Marchetti & Nevitt, 2003](https://www.researchgate.net/publication/226487156_Effects_of_Hatchery_Rearing_on_Brain_Structures_of_Rainbow_Trout_Oncorhynchus_mykiss) | [Shigeno et al. 2018](https://www.frontiersin.org/articles/10.3389/fphys.2018.00952/full?source=post_page-----b9bac630960b----------------------) |  |  |  | [Sayol et al. 2020](https://royalsocietypublishing.org/doi/full/10.1098/rspb.2020.0762) |  |
| Brain mass to body mass ratio | [Minervini et al., 2016](https://journals.plos.org/plosone/article?id=10.1371/journal.pone.0157378) | [Olkowicz et al. 2016](https://www.pnas.org/content/113/26/7255) | [Kotrschal & Palzenberger 1992](https://link.springer.com/article/10.1007/BF00002560) | [Wiper et al., 2014](https://cdnsciencepub.com/doi/abs/10.1139/cjfas-2013-0624) | [Packard 1971](https://onlinelibrary.wiley.com/doi/epdf/10.1111/j.1469-185X.1972.tb00975.x) |  |  |  | [Gowda & Gronenberg, 2019](https://arizona.pure.elsevier.com/en/publications/brain-composition-and-scaling-in-social-bee-species-differing-in-) |  |
| Brain volume | [Kruska and Röhrs, 1974](https://pubmed.ncbi.nlm.nih.gov/4851103/) | [Henriksen et al. 2016](https://www.nature.com/articles/srep34031) | [Kotrschal et al. 1991](https://link.springer.com/chapter/10.1007/978-94-011-3092-9_10) | [Peris Tamayo et al. 2020](https://onlinelibrary.wiley.com/doi/full/10.1002/ece3.6771) |  |  | [Krieger et al. 2012](https://link.springer.com/article/10.1007/s00441-012-1353-4) |  | [Gowda & Gronenberg, 2019](https://arizona.pure.elsevier.com/en/publications/brain-composition-and-scaling-in-social-bee-species-differing-in-) | [Snell-Rood et al. 2020](https://onlinelibrary.wiley.com/doi/epdf/10.1111/evo.14072?saml_referrer) |
| Critical flicker-fusion frequency in Hz | [Rosolen et al. 2005](https://iovs.arvojournals.org/article.aspx?articleid=2401734) | [Lisney et al. 2011](https://pubmed.ncbi.nlm.nih.gov/21527269/) | [Hanyu & Ali 1964](https://onlinelibrary.wiley.com/doi/abs/10.1002/jcp.1030630306) | [Hanyu & Ali 1964](https://onlinelibrary.wiley.com/doi/abs/10.1002/jcp.1030630306) | [Bullock & Budelmann 1991](https://link.springer.com/article/10.1007/BF00217112) | [Kingston et al. 2020](https://royalsocietypublishing.org/doi/10.1098/rsbl.2020.0298) | [Frank et al. 2012](https://journals.biologists.com/jeb/article/215/19/3344/10962/Light-and-vision-in-the-deep-sea-benthos-II-Vision) | [Waterman 1961](https://books.google.co.uk/books?hl=en&lr=&id=oOlO3lCeR0wC&oi=fnd&pg=PA1&ots=AGwHUcAbYx&sig=tAEHYNv00309iI7E1BwmVzTm0_c&redir_esc=y#v=onepage&q&f=false) | [Meyer-Rochow, 2019](http://journal.bee.or.kr/xml/20517/20517.pdf) |  |
| Encephalization quotient | [Minervini et al., 2016](https://journals.plos.org/plosone/article?id=10.1371/journal.pone.0157378) | [Kumar Panigrahy et al. 2017](https://www.researchgate.net/publication/337589479_GROSS_MORPHOLOGY_AND_ENCEPHALIZATION_QUOTIENT_OF_BRAIN_IN_MALE_AND_FEMALE_VANARAJA_CHICKENS_AT_DIFFERENT_AGES) | [Masai et al. 1983](https://doi.org/10.1111/j.1439-0469.1983.tb00256.x) | [Triki et al. 2021](https://doi.org/10.1159/000520741) |  |  |  |  |  |  |
| Neuron packing density | [Friede 1954](https://www.karger.com/Article/Abstract/140905) | [Olkowicz et al. 2016](https://www.pnas.org/content/113/26/7255) |  |  |  |  |  |  |  |  |
| Total neurons | [Jelsing et al., 2006](https://pubmed.ncbi.nlm.nih.gov/16574805/) | [Olkowicz et al. 2016](https://www.pnas.org/content/113/26/7255) | [Zupanc et al. 2005](https://doi.org/10.1002/cne.20571) | [Zupanc et al. 2005](https://doi.org/10.1002/cne.20571) |  |  | [Feinberg & Mallatt 2016.](https://books.google.ca/books?hl=en&lr=&id=fIPTCwAAQBAJ&oi=fnd&pg=PR5&dq=Todd+Ancient+origin+of+consciousness&ots=c6jVYS29q0&sig=rvzDnoIVFK5AxAz2XaLmoyZW_94#v=onepage&q=Todd%20Ancient%20origin%20of%20consciousness&f=true) | [Kondoh and Hisada 1986](https://onlinelibrary.wiley.com/doi/abs/10.1002/cne.902540209) | [Gowda & Gronenberg, 2019](https://arizona.pure.elsevier.com/en/publications/brain-composition-and-scaling-in-social-bee-species-differing-in-) | [Fukushima & Kanzaki 2009](https://pubmed.ncbi.nlm.nih.gov/19148932/) |
| Working memory load | [Arts et al., 2009](https://www.sciencedirect.com/science/article/abs/pii/S0166432809003696) | [Nordquist et al. 2011](https://www.researchgate.net/publication/229052519_Laying_hens_selected_for_low_mortality_Behaviour_in_tests_of_fearfulness_anxiety_and_cognition) | [Bloch et al. 2019](https://www.sciencedirect.com/science/article/pii/S0166432819300385) | [Sovrano et al. 2018](https://www.frontiersin.org/articles/10.3389/fpsyg.2018.02341/full) | Borrelli & Fiorito 2008 |  |  | [Tierney and Andrews, 2013](https://link.springer.com/article/10.1007/s10071-012-0547-1) | [Siviter et al., 201](https://besjournals.onlinelibrary.wiley.com/doi/abs/10.1111/1365-2664.13193)8 | [Blakiston et al. 2008](https://journals.plos.org/plosone/article?id=10.1371/journal.pone.0001736) |
